# Thyroid Hormone T4 Mitigates Traumatic Brain Injury by Dynamically Remodeling Cell Type Specific Genes, Pathways, and Networks in Hippocampus and Frontal Cortex

**DOI:** 10.1101/2022.11.15.516648

**Authors:** Guanglin Zhang, Graciel Diamante, In Sook Ahn, Victoria Palafox-Sanchez, Jenny Cheng, Michael Cheng, Zhe Ying, Susanna Sue-Ming Wang, Kevin Daniel Abuhanna, Nguyen Phi, Douglas Arneson, Ingrid Cely, Kayla Arellano, Ning Wang, Fernando Gomez-Pinilla, Xia Yang

## Abstract

**Background:** The complex pathology of mild traumatic brain injury (mTBI) is a main contributor to the difficulties in achieving a successful therapeutic regimen. Thyroxine (T4) administration has been shown to prevent the cognitive impairments induced by mTBI in mice.

**Method:** To understand the underlying mechanism, we carried out a single cell transcriptomic study to investigate the spatiotemporal effects of T4 on individual cell types in the hippocampus and frontal cortex at three post-injury stages.

**Findings:** Our multi-tissue multi-stage results showed that T4 treatment altered the proportions and transcriptomes of numerous cell types across tissues and timepoints, particularly oligodendrocytes, astrocytes, and microglia, which are crucial for injury repair. T4 also reversed the expression mTBI-affected genes such as *Ttr, mt-Rnr2*, *Ggn12, Malat1, Gnaq,* and *Myo3a*, as well as numerous pathways such as cell/energy/iron metabolism, immune response, nervous system, and cytoskeleton-related pathways. Cell-type specific network modeling revealed that T4 mitigated select mTBI-perturbed dynamic shifts in subnetworks related to cell cycle, stress response, and RNA processing in oligodendrocytes. Cross cell-type ligand-receptor networks recapitulated the roles of App, Hmgb1, Fn1, and Tnf in mTBI, the latter two ligands having been previously identified as TBI network hubs. mTBI and/or T4 signature genes were enriched for human genome-wide association study (GWAS) candidate genes for cognitive, psychiatric and neurodegenerative disorders related to mTBI, supporting T4 as a potential mTBI treatment.

**Interpretation:** Our systems-level approach elucidated the temporal and spatial dynamic reprogramming of cell-type specific genes, pathways, and networks, as well as cell-cell communications through which T4 mitigates cognitive dysfunction induced by mTBI.

**Funding:** This work was funded by NIHR01NS117148 to X.Y. and F.G.P.

**Research in Context:** *Evidence before this study:* Dysfunction in the brain resulting from traumatic brain injury can display immediately as well as several years post-injury. It also impacts various brain regions, including the hippocampus and frontal cortex, which are linked to distinct disease pathologies. The complexity of spatiotemporal and molecular dynamics of perturbation caused by TBI hinder our ability to establish an effective therapeutic approach. Recently, thyroid hormone poses promise as a potential therapeutic target based on our previous scRNA-seq studies. Yet, the mechanisms by which T4 alleviates mTBI, specifically those related to spatial, temporal, and cell-type specificity, remain unexplored.

*Added value of this study:* We examined the impact of T4 intervention in mitigating mTBI by investigating the transcriptome and functional pathways across two affected brain regions, the frontal cortex and hippocampus, in different stages of injury. Utilizing a systems biology approach, we conducted within- and between-cell-type network modeling, cell-cell communication and integrating human genome-wide association studies (GWAS) analysis. This comprehensive strategy aimed to elucidate the cellular and molecular mechanisms through which T4 averts cognitive impairments induced by mTBI.

*Implications of all the available evidence:* Our findings offer molecular evidence that the administration of T4 impacts a wide range of genes, biological processes, and networks, thereby preventing the advancement of mTBI-induced brain dysfunction and associated diseases. This comprehensive impact of T4 suggests potential advantages in efficacy compared to other therapeutic options that concentrate on specific pathways and targets.

## Introduction

Traumatic brain injury (TBI) has an estimated incidence of 69 million per year and is a primary cause of injury-induced death worldwide.^1^ Based on statistics from Centers for Disease Control and Prevention, approximately 61,000 TBI-related deaths were recorded in the United States in 2019, averaging 166 deaths per day.^2^ Mild TBI (mTBI) accounts for approximately 81% of TBI cases globally, affecting ∼55·9 million people each year.^1^ Recent studies have shown mTBI exhibit distinct pathophysiology from moderate/severe TBI, representing a different disease state and therefore requiring its own treatment regimen.^3–5^

TBI/mTBI disease pathology has both a temporal and spatial component, in that TBI-induced brain dysfunction can start acutely and persist for many years after injury and affect multiple brain regions such as the hippocampus and frontal cortex, which are associated with different disease pathologies.^6–9^ In addition, numerous molecular pathways have been implicated in the pathogenesis of TBI/mTBI, including excitotoxicity, mitochondrial dysfunction, oxidative stress, lipid peroxidation, neuroinflammation, axon degeneration, and apoptotic cell death.^8–13^

The broad spectrum of spatial, temporal, and molecular dysfunctions associated with TBI hamper our ability to achieve a successful therapeutic regimen. In particular, current treatments for moderate/severe TBI such as glutamate receptor antagonists, inhibitors of calcium-related signals, antioxidants, anti-inflammatory drugs, anti-apoptotic agents, and neurotrophic factors, as well as the targeting of reactive oxygen species using methylprednisolone for mTBI, have yielded limited degrees of success.^12^

Multiple recent lines of evidence support that the thyroid hormone pathway may be a more effective therapeutic avenue which can target a broad spectrum of molecular pathways involved in TBI/mTBI. Thyroid hormones such as thyroxine (T4) and triiodothyronine (T3) are essential for normal brain development and function and have roles in diverse physiological processes such as temperature regulation, energy balance and metabolism. Previous studies have shown reduced T4 and T3 levels in patients who suffered head trauma and that these levels were negatively correlated with injury severity and poor outcome.^14^ Post-TBI administration of T3 has been shown to significantly improve motor and cognitive recovery, reduce lesion volume, inhibit neuroinflammation, and induce the expression of neuroprotective neurotrophins BDNF and GDNF.^15^ Additionally, our previous studies using single-cell RNA sequencing (scRNAseq) to investigate the cell-type specific dynamics of mTBI revealed transthyretin (*Ttr*), a T4 transporter, as a potential therapeutic target.^8^ *Ttr* transports T4 to the brain, where it is converted into the active hormone T3, which can bind and activate thyroid hormone receptors to regulate various brain functions. We have shown that the T4 treatment significantly reversed not only the cognitive impairment caused by mTBI but also mTBI-induced perturbations in metabolic pathways based on hippocampal bulk RNA sequencing analysis.^8^ However, the spatial, temporal and cell type specific mechanisms through which T4 mitigates mTBI have yet to be investigated.

Here, we focused on investigating the transcriptome and functional pathways and networks following T4 intervention in mTBI using scRNAseq on two brain regions impacted by mTBI, the frontal cortex and hippocampus, at acute (24hr), subacute (7-day), and subchronic (21-day) stages. We applied a systems biology approach involving within- and between-cell-type network modeling and the integration of human genome-wide association studies (GWAS) to understand the cellular and molecular aspects of how T4 prevents mTBI-induced cognitive impairments (Fig.1a).

**Fig. 1.**
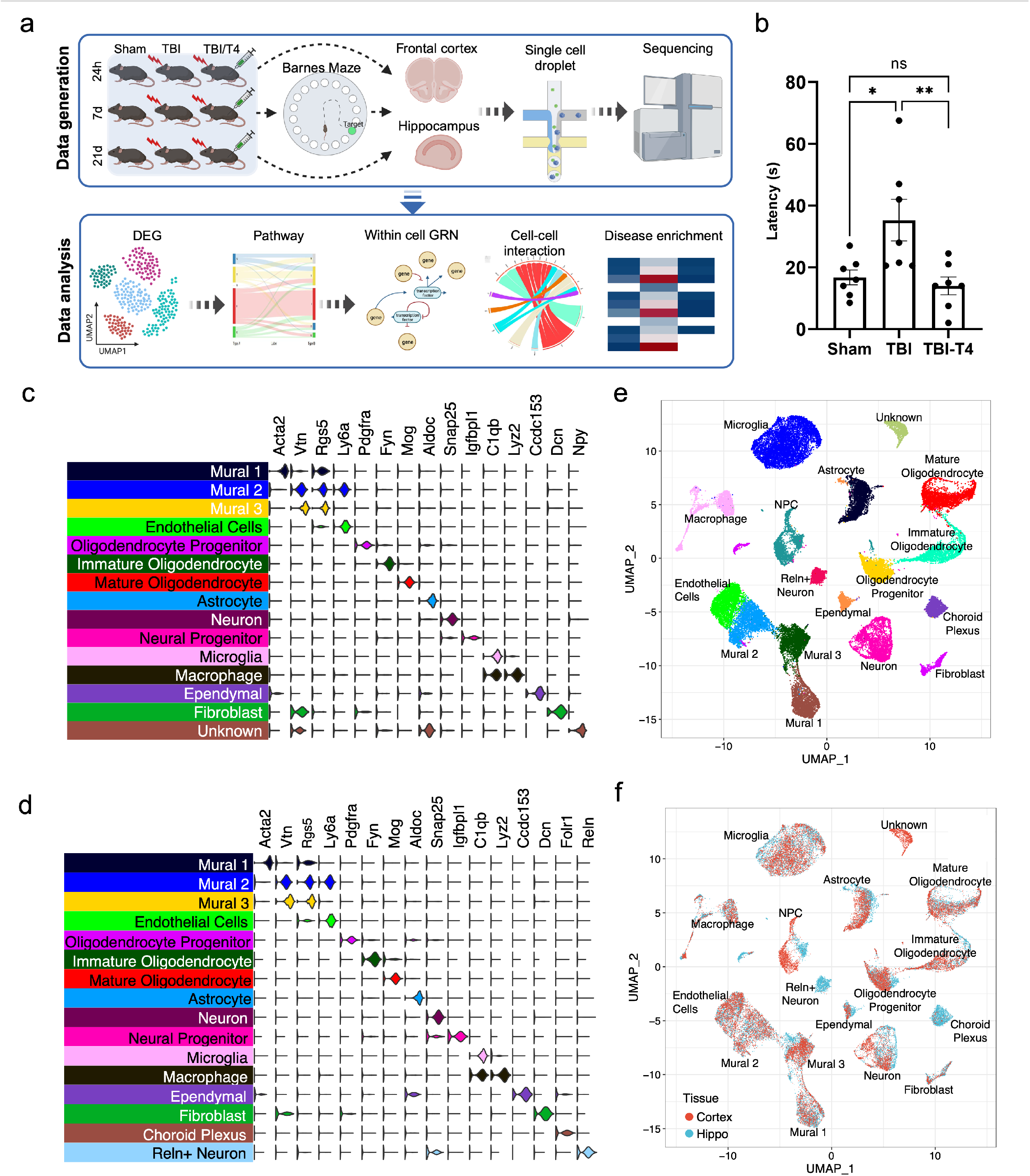
Overall study design of the T4 treatment study, Barnes Maze memory test, and cell type clustering from scRNAseq across two tissues. a) Schematic of study design. Nine groups of mice (n=7/group) including sham, mTBI, and mTBI+T4 treatments across 24hr, 7-day, and 21-day timepoints were investigated. Mice from the 7-day timepoint underwent spatial learning and memory evaluation using the Barnes Maze. The hippocampus and frontal cortex tissues of a subset of mice from each of the 9 groups (n=3/group) were then subject to scRNAseq analysis. scRNAseq data was subsequently analyzed to identify the differentially expressed genes and perturbed pathways and networks in individual cell types, altered cell-cell communications, and human disease relevance of the TBI effect and T4 treatment effect at individual timepoints and in each brain region. b) Barnes Maze memory test for mice from the 7-day timepoint. T4 treatment reversed the TBI-induced increase in latency time for mice to find the escape hole. Longer latency time indicates compromised memory function. Statistics were computed using one-way ANOVA with Tukey correction for multiple comparisons. *p < 0·05, **p < 0·01, ns = not significant, n = 7 per group. C and D) Distinct expression of unique marker genes for each cell type in the frontal cortex (c) and hippocampus (d). e and f) UMAP embeddings of cells according to cell types (e) and tissues (f; frontal cortex vs. hippocampus). Each point in e and f represents a single cell. Cells were clustered based on transcriptome similarity using Louvain clustering, and cell types were labeled based on the expression of canonical cell type markers in (c) and (d). Within each tissue n=3 independent animals per group were used for scRNAseq.

## Methods

### Animals and fluid percussion injury (FPI) model of mTBI

Ten-week-old male C57BL/6J (B6) mice from Jackson Laboratory weighing between 20 and 25g were maintained under standard housing conditions (room temperature 22-24 °C) with 12 hours (h) light/dark cycle. Mice were randomly divided into either the sham (7 mice per timepoint) or mTBI group (14 mice per timepoint), and mTBI mice were further divided into two groups with or without T4 treatment (7 mice with TBI only, 7 mice with TBI plus T4 per timepoint). Brain injury was induced using a fluid percussion injury technique. Craniotomy was performed under a microscope (Leica), where a 3·0-mm diameter hole was made 2·5mm posterior to the bregma and 2·0mm lateral (left) of the midline with a high-speed drill (Dremel, Racine, WI, USA). The animals with intact dura underwent the fluid percussion injury procedure. Specifically, a plastic cap was first fixed over the hole with adhesive and dental cement. When the dental cement hardened, the cap was filled with 0.9% saline solution. Anesthesia was discontinued, and the injury cap was connected to the fluid percussion device. A fluid percussion (1·5–1·7 atm) was administered and wake-up time was determined to fall into the range of 5-12 min. The animals in the sham group also underwent an identical preparation except for the percussion. T4 (1·2 ug/100g bodyweight) or saline (control) was administered via i.p. injection at 1hr and 6hr post-FPI, based on previous studies where a therapeutic effect was observed.^8,9,16^ Mice were sacrificed and tissue collection was conducted at 24hr, 7-day and 21-day post treatment. Hippocampus and frontal cortex from each mouse were used for scRNAseq using the Drop-seq method.

### Ethics

This study was performed following the National Institutes of Health Guide for the Care and Use of Laboratory Animals, USA. The experimental protocol was approved by the Chancellor’s Animal Research Committee of the University of California at Los Angeles.

### Behavioral assessments

The memory retention of mice from Sham, TBI, and T4 treatment groups were assessed using the Barnes Maze test seven days post-injury (n=7/group), as previously described.^8,9^ The learning phase involved training animals with two trials per day over four consecutive days, followed by a test of memory retention two days after the last learning session.

The Barnes maze used in this study was constructed from 1·5 cm thick acrylic plastic, measuring 120 cm in diameter, and featured 40 evenly spaced holes, each 5 cm in diameter, along its edges. The maze was well-lit with 900 lumens from four overhead halogen lamps, creating an aversive environment that prompted the mice to search for a concealed dark escape chamber located beneath one of the holes along the maze’s perimeter.

To monitor the trials, a video camera was positioned directly overhead at the center of the maze, recording all test sessions simultaneously. Each trial began with the placement of the mouse in the center of the maze, covered by a cylindrical starting chamber. After a 10-second delay, the starting chamber was lifted. The training sessions ended either when the animal successfully entered the escape chamber or when a pre-determined time limit of 5 minutes was reached, whichever came first. To prevent potential interference from previous animals, all surfaces were routinely cleaned before and after each trial, ensuring the elimination of any olfactory cues.

### Single-cell isolation for scRNAseq

For single cell profiling, tissues from a subset of mice from each of the 9 groups (n=3/group/tissue) were subject to scRNAseq analysis for a total of 54 samples. One cortex sample from 21-day T4 group didn’t pass QC and was removed from downstream analysis. Single-cell suspensions were prepared as previously described.^8,9,17^ Briefly, the hippocampus and frontal cortex were dissected from the ipsilateral side of the brain, sliced into thin pieces, and transferred into 4□ml HABG medium containing Hibernate A, B27 supplement, and 0·5 mM Glutamax (Fisher Scientific, Hampton, NH, USA) then incubated in a water bath at 30□°C for 8□mins. Subsequently, the HABG media was removed and prewarmed papain solution (2□mg/ml HA-Ca) was added to the tissues and incubated for 30 mins at 30□°C with shaking. The papain solution was then washed out with 2 ml prewarmed HABG. The tissue was triturated approximately ten times for 45s using a siliconized 9-in Pasteur pipette with a fire-polished tip. After 1 min, the supernatant was gently transferred to the top of the prepared OptiPrep density gradient (Sigma Aldrich, St. Louis, MO, USA) without breaking the gradient. After centrifugation of 800□g for 15□min at 22□°C, the top 6□ml containing cellular debris was discarded. The bottom fraction was collected and diluted with 5 ml HABG and centrifuged for 3□min at 22□°C at 200□g followed by removal of the supernatant containing the debris. Finally, the cell pellet was loosened by flicking the tube and re-suspended in 1□ml of 0·01% BSA in PBS. The cell suspension was filtered through a 40-micron strainer (Fisher Scientific, Hampton, NH, USA) followed by cell counting.

### Single cell barcoding and library preparation

Drop-seq was performed as previously described, using protocol version 3·1.^18^ The single-cell suspensions were prepared at a final concentration of 100 cells/μl. The cell suspension, oil (EvaGreen, Bio-Rad, Hercules, CA, USA), and lysis buffer mixed with barcoded beads (ChemGenes, Wilmington, MA, USA) were co-flowed through a microfluidic device (FlowJEM, Toronto, Canada) at fixed speeds (oil: 15,000 μl/h, cells: 4000 μl/h, and beads 4000 μl/h) to generate droplets. We dispensed 4000 beads into each PCR tube and ran 4+11 cycles. Pooled PCR tubes proceeded to cDNA library preparation. The cDNA library quality was checked using a BioAnalyzer high-sensitivity chip (Agilent, Santa Clara, CA, USA). The cDNA library was then fragmented and indexed using the Nextera DNA Library Preparation kit (Illumina, San Diego, CA, USA). The library quality was checked on a BioAnalyzer high-sensitivity chip and concentration was quantified by Qubit assays (ThermoFisher, Canoga Park, CA, USA) before sequencing.

### Sequencing of single cell libraries

Sequencing was performed on an Illumina HiSeq 4000 (Illumina, San Diego, CA, USA) using the Drop-seq custom read 1B primer (GCC TGT CCG CGG AAG CAG TGG TAT CAA CGC AGA GTA C) (IDT, Coralville, IA, USA) and paired-end 100bp reads were generated. Read 1 consists of the 12 bp cell barcode, followed by the 8 bp unique molecular identifier (UMI), and the last 80 bp on the read are not used. Read 2 contains the single-cell transcripts.

### Single cell data pre□processing and quality control

The fastq files of the Drop-seq sequencing data were processed to digital expression gene matrices using Drop-seq tools version 1·13 (https://github.com/broadinstitute/Drop-seq) and dropEst as previously described.^9^ We followed a modified version of the snakemake-based dropSeqPipe (https://github.com/Hoohm/dropSeqPipe) workflow as previously described.^9^ Briefly, reads with low-quality barcodes were removed, and the cleaned reads were aligned to the mouse reference genome mm10 using STAR-2·5·0c. The reads which overlapped with exons were tagged using a RefFlat annotation file of mm10. The Drop-seq Tools function DetectBeadSynthesisErrors was used to estimate a bead synthesis error rate of 5–10%, within the acceptable range. Finally, we generated a digital gene expression matrix for each sample where each row is the read count of a gene, and each column is a unique single cell. The transcript counts of each cell were normalized by the total number of UMIs for that cell. These values are then multiplied by 10,000 and log-transformed. Single cells were identified from background noise using a threshold of at least 200 genes and 300 transcripts. To further control the data quality, cells containing gene numbers between 200 and 3000, and that had mitochondrial gene content of <15% were used for analysis.

### Identification of cell clusters

After quality control, the Seurat V4 package^19^ was used for dimension reduction and the Louvain algorithm^20^ was used to cluster the cells. All cells were then projected onto two dimensions using uniform manifold approximation and projection (UMAP). Cells with similar transcriptional expression patterns are plotted closer together forming a cluster. Canonical correlation analysis (CCA)^21^ was used to integrate and align data across different timepoints or conditions. The optimal cluster number was determined using Jackstraw permutation.

### Identification of marker genes and cluster cell types

Cell cluster-specific marker genes were identified using the FindConservedMarkers function in Seurat. Briefly, the Wilcoxon Rank Sum Test was performed within each set of samples and a meta p-value across all conditions was computed to assess the significance of each gene as a marker for a cluster across datasets of different timepoints and conditions. To be considered in the analysis, a gene had to be expressed in at least 10% of the single cells from at least one of the groups. Multiple testing was corrected using the Bonferroni method on the meta p-values. The genes with an adjusted p-value < 0·05 were defined as cell-type-specific marker genes. To determine cell types, cell cluster-specific marker genes were compared against known cell marker genes for hippocampus^22^ and frontal cortex^23^ as well as single cell annotated datasets from DropVIZ atlas^24^ as previously described.^8,9^

### Euclidean Distance analysis to determine global shifts in each cell cluster between treatments

To quantify the effect of TBI and T4 treatment on each cell type, average gene expression profiles were calculated across cells within each group and cell type, such that each treatment group contains a representative cell for each cell type. The global transcriptomic shift for each cell type was then measured using Euclidean distance between the representative cells for the mTBI vs. sham and T4 vs. mTBI group comparisons. Due to the high gene expression variation, we selected the top 1000 expressed genes and standardized their expression. To determine significance of the transcriptomic effect in each comparison, we generated a null distribution of Euclidean distances for each cell type by measuring the distance between groups of randomly sampled cells from the given cell type of the two groups being compared. After 1000 permutations, we compared the Euclidean distance between the real treatment groups and the null distribution to determine the significance. Bonferroni correction was used for multiple testing corrections across cell types.

### Identification of differentially expressed genes (DEGs) and pathway annotation

DEG analysis between T4 treatment and TBI groups as well as between TBI and sham control samples within each identified cell type at each time point was done using a Wilcoxon Rank Sum Test. For DEG analysis, we considered only the genes which were expressed in at least 10% of the cells from one of the two groups in a given comparison. DEGs were defined as genes with a Bonferroni corrected adjusted p-value < 0·05. The DEGs were then used in pathway analysis, where enrichment of pathways from KEGG, Reactome, BIOCARTA, GO Molecular Functions, and GO Biological Processes were assessed using Fisher’s exact test. Significant pathways were defined with the Benjamini–Hochberg corrected false discovery rate (FDR) < 0·05.

### Within-cell-type gene network analysis

Within-cell-type gene regulatory networks (GRNs) were inferred in each tissue for each cell type across all timepoints and treatments. We used SCING^25^ to construct the GRNs, which uses a bootstrapped gradient boosting regression to infer robust gene interactions. Subnetworks, or modules, were detected in each cell type GRN using the Leiden clustering algorithm, and the AUCell scoring method from SCENIC^26^ measured module activity in each cell based on ranked expression of module genes. We annotated the modules with enriched pathways using the enrichR R API.^27^ To predict the effect of TBI and T4 on module activity, we used the lm() function in R to create a linear regression model to predict module expression from treatment, time point, and their interactions. We treated each variable as categorical and interpreted their respective coefficients as the variable’s effect on module expression relative to the corresponding control group (i.e. sham as control for treatment groups or 24hr as control for timepoint effect). We controlled the FDR using Benjamini Hochberg correction. Network visualizations were performed using Cytoscape.^28^

### Between-cell-type communication analysis

Ligand-target gene connections were predicted in each tissue type and at each timepoint with Nichenet, which combined cell expression data with prior knowledge on ligand-target signaling paths, signal transduction, and gene regulatory interactions.^29^ Astrocytes, endothelial cells, mature oligodendrocytes, neurons, and NPCs were defined as the receiver/target cell populations, whereas all cell types were defined as the sender populations. Significant DEGs (adjusted p-value < 0·05) between mTBI vs. sham and T4 vs. mTBI for the target cell populations were used to predict the corresponding upstream ligands and intercellular communications. Genes consistently predicted as ligands at every timepoint for each tissue were further examined for differential expression patterns in mTBI vs. sham and T4 vs. mTBI. A cell type was considered a sender of a particular ligand if the expression of the ligand was greater than the sum of the average expression of the ligand and half of one standard deviation of all the cell types. A maximum of five DEGs, based on adjusted p-value, was visualized for each comparison and cell type in each circle plot.

### Disease association analysis

To assess whether our mouse mTBI- and T4-induced gene signatures are associated with human diseases and traits, we collected candidate genes from the GWAS catalog database^30^ and then determined whether our cell type specific DEGs were enriched for human disease/trait genes. The DEGs used for the enrichment analysis had a P-value <0·01. Enrichment of disease/trait associated genes was based on a hypergeometric test followed by multiple testing corrections with the Benjamini-Hochberg method. Disease/trait gene enrichment that was considered significant had an FDR < 0·05 and an overlap gene of >= 3 genes.

## Results

### T4 treatment reverses memory impairment caused by brain injury

We carried out sham surgery, fluid percussion injury (FPI) at 1·5–1·7 atm as a form of mTBI, and FPI followed by T4 injection intraperitoneally at 1·2 ug/100g bodyweight dosage twice at 1hr and 6hr post-injury, as done previously, where a protective effect on cognition was observed.^8^ The sham and mTBI groups were also injected with saline vehicle as the control for T4 treatment. Subsets of mice from each treatment group were kept for 24hr, 7-day, and 21-day (n=7/group/timepoint) before further experimentation.

To test the spatial learning and memory affected by mTBI and mTBI+T4 treatments, a Barnes maze test was applied 7 days after injury.^31^ Memory testing was not carried out at other timepoints because mice at 24hr post-injury were not capable of performing memory tests and previous studies have shown that mice at 21-day post-injury achieved cognitive recovery from mTBI^32^. T4 treatment significantly improved the memory performance at 7-day post-injury as revealed by the restoration of the latency time to a level that closely matched that of the sham group at the subacute phase (Fig.1b), confirming our previous results.^8^

### Identification of cell types from the frontal cortex and hippocampus

To understand the molecular mechanisms of T4 treatment, we carried out scRNAseq on the hippocampus and frontal cortex from each treatment group at each timepoint (n=3/group/timepoint). Across 54 samples from three groups of mice at three timepoints, we sequenced 133,523 single cells that passed quality control (Supplementary Fig.S1). One cortex sample didn’t pass quality control and was removed from downstream analysis. Cell clusters were identified across tissues, and cluster identities were determined using canonical marker genes identified from the previous single cell studies in atlas.^22,23,24^ Among these clusters, 14 general cell types were identified in the cortex (Fig.1c) and 16 in the hippocampus (Fig.1d) based on the distinct expression of known marker genes in individual cell types in the two brain regions. There were 14 general cell types that were concordant between the hippocampus and cortex (Fig.1c-1f), but the choroid plexus and Rel+ neuronal clusters were unique to the hippocampus.

We further extracted the neuronal cluster and carried out subclustering analysis to identify neuronal subtypes, revealing hippocampal neuronal subtypes such as CA1/CA3 pyramidal neurons and dentate gyrus (DG) granular cells, and frontal cortex neuronal subtypes such as L2/3 and L6 neurons, various interneurons, and GABA1/GABA2 neurons (Supplementary Fig.S2a and S2b).

These results support that our scRNAseq studies successfully retrieved known cell types and subtypes in each of the brain regions examined.

### Spatiotemporal shifts of cell proportions following T4 treatment across timepoints

We first assessed the cell proportion changes affected by mTBI and T4 treatments. As shown in Figure 2, mTBI induced changes in the proportions of numerous cell types in the hippocampus at 24hr, both hippocampal and cortex cell types at 7-day, but fewer changes at 21-day post-injury. In contrast, T4 treatment affected the proportions of a smaller number of cell types, many of which had the same direction of change as mTBI compared to sham animals. However, we note that T4 reversed the mTBI-induced decrease of the hippocampal mural 3 population at 7-day post-mTBI (labeled with red asterisk in Fig.2b) as well as the mTBI-induced increase in the hippocampal endothelial cell population at 21-day post-mTBI to similar levels as the sham group (labeled with red asterisks in Fig.2c). Additionally, in the hippocampus, T4 treatment uniquely increased endothelial cells at 7-days and increased the choroid plexus population at 21-days (labeled with cyan asterisks in Fig.2b-c). Notably, choroid plexus is not part of the hippocampus but can easily be included during hippocampus dissection.^33^ Endothelial and mural cells are important for blood-brain-barrier (BBB) integrity and function while choroid plexus plays a role in neural repair.^34^ Therefore, T4 treatment altered the proportions of multiple cell types in or near the hippocampus that either counteracted mTBI-induced changes or enhanced protective and reparative cell types in response to injury. In contrast, T4-specific cell proportion changes in the frontal cortex were only found for a mural subtype.

**Fig. 2.**
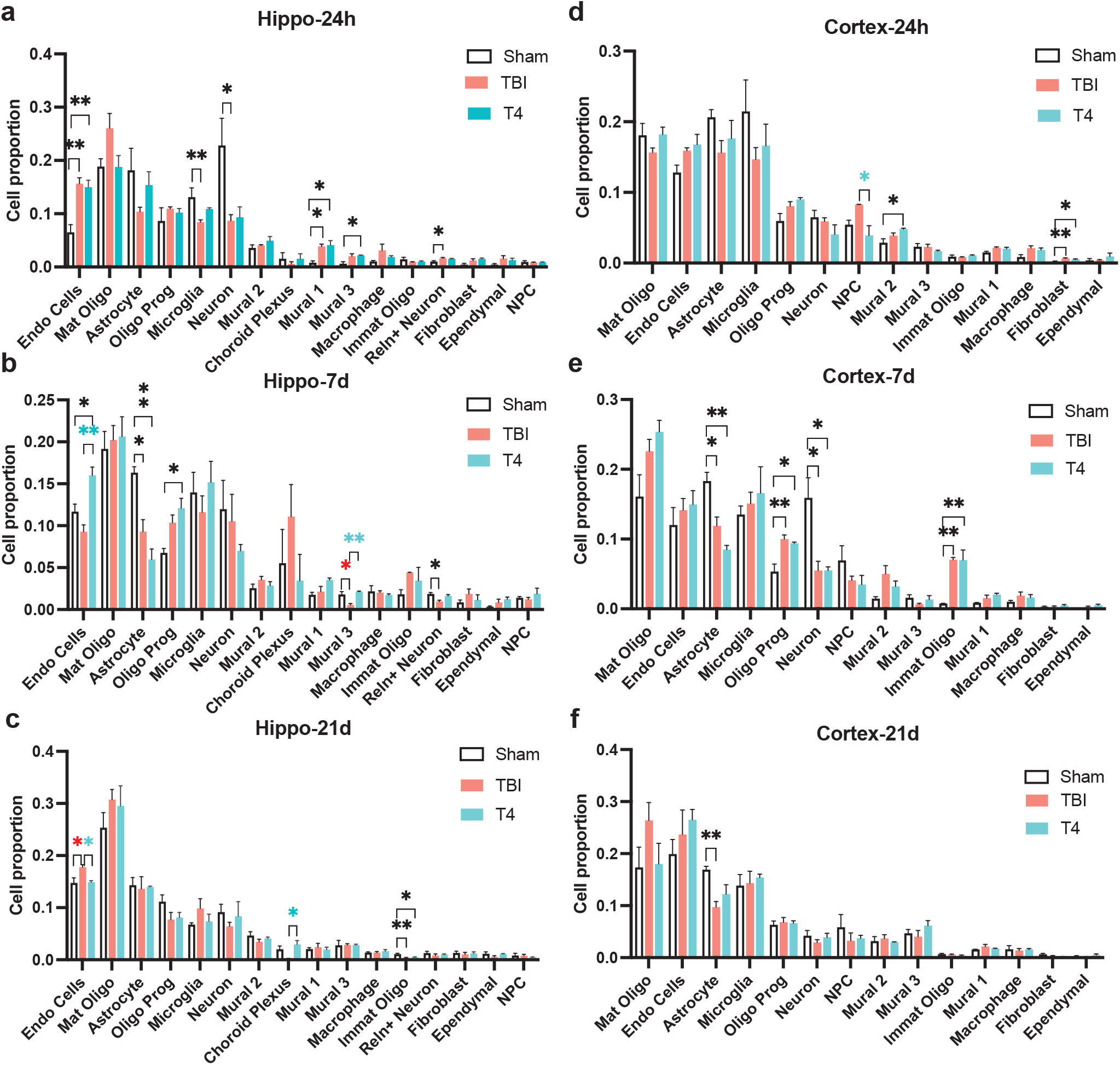
Spatiotemporal changes of cell proportions in response to mTBI or T4 treatments. Statistics were computed using one-way ANOVA with Tukey correction for multiple comparisons. *p < 0·05, **p < 0·01, n = 3 per group/timepoint. Pairs of contrasting red and cyan asterisks denote the opposite change directions between mTBI (red) and T4 treatments (cyan). Cell types uniquely altered in T4 treatment are annotated with cyan asterisks.

### Spatiotemporal shifts of the global transcriptome following mTBI and T4 treatments

Even without cell proportion changes, shifts in the gene expression profiles can also signify sensitivity or vulnerability of a cell type to injury or treatments. Indeed, visualization of the cell clusters by tissue, timepoint, and treatment groups revealed subtle shifts in the cells between groups in each brain region at each timepoint (Fig.3a; frontal cortex plots in top row, hippocampus plots in bottom row; time points in columns). To quantify the global transcriptomic shifts within each cell type between treatment groups, we used a Euclidean distance based method,^8^ which measures the distance between two treatment groups in the gene expression profiles represented by the top 1000 highly expressed genes in each cell type. A larger Euclidean distance is an indicator of stronger transcriptomic responses to treatment. Most cell types in both the cortex and hippocampus exhibited transcriptomic shifts in response to mTBI (mTBI vs. sham) but not T4 treatment (T4 vs. mTBI) at the acute (24hr) and subacute (7-day) stages (Fig.3a top/middle panels; red dots). At the subchronic 21-day post-TBI, only a few cell types showed significant global transcriptomic response to mTBI in the hippocampus (Fig.3b bottom panels). In terms of T4 treatment effect on the global transcriptome, significant changes only occurred at and after the 7-day stage. In the frontal cortex, mature oligodendrocytes (7-day) and neurons and neural progenitor cells (21-day) were responsive to T4 (Fig.3b left middle/bottom panels, cyan dots); in the hippocampus, endothelial cells (7-day), astrocytes (21-day), and mature oligodendrocytes (7-day and 21-day) responded to T4 treatment (Fig.3b right middle/bottom panels). These results indicate that mTBI had dramatic effects on the global transcriptome across cell types and tissues at the acute and subacute phases, whereas T4 treatment mainly caused global transcriptomic shifts in a select set of cell types from mTBI at the subacute and subchronic phase, suggesting a slower and more selective effect of T4 treatment compared to the mTBI effect.

**Fig. 3.**
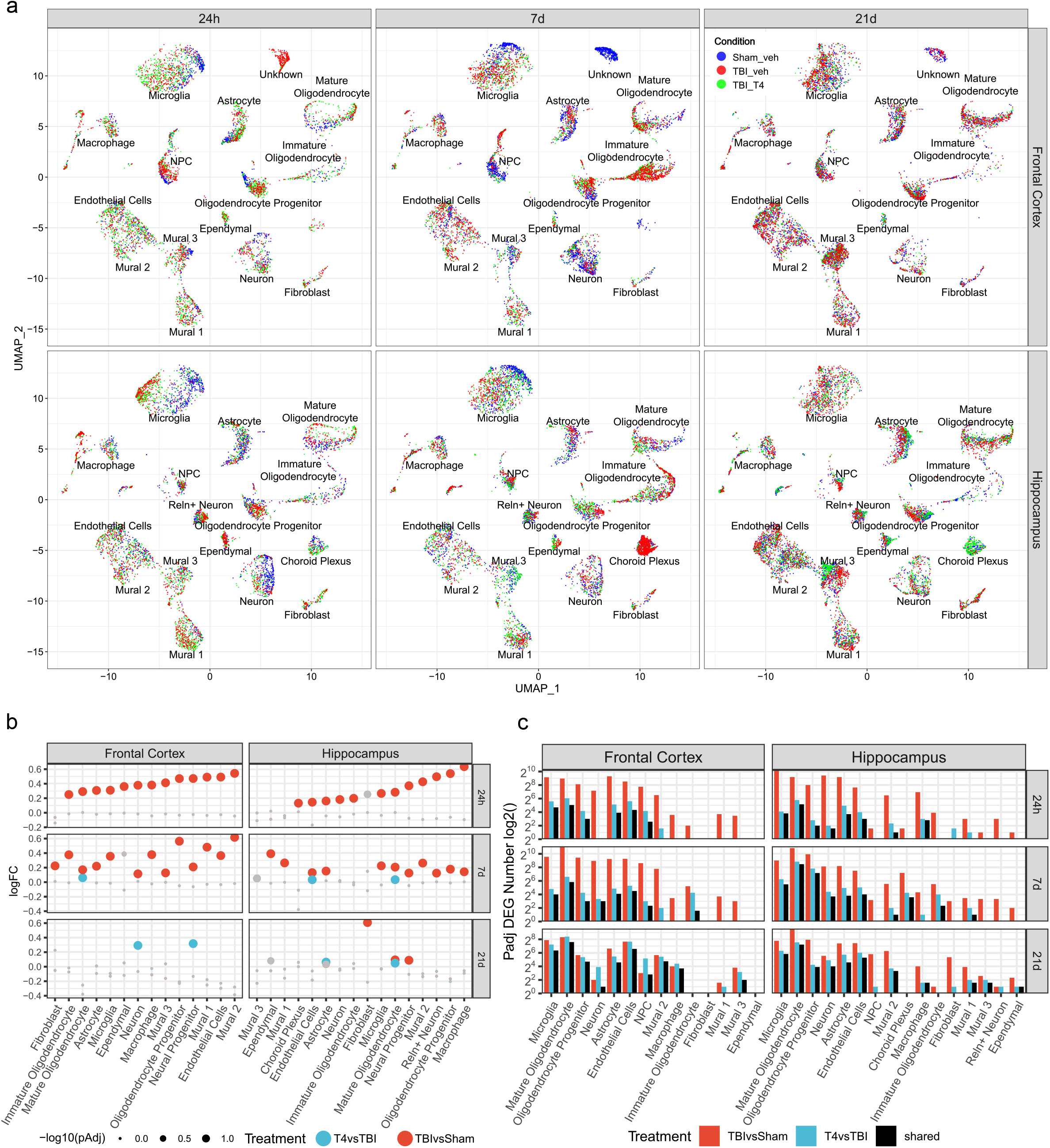
mTBI and T4 treatments induced transcriptomic shifts across cell types in the frontal cortex and hippocampus. (a) UMAP embeddings showing cells colored by treatment groups (blue for sham, red for TBI, green for T4) within individual cell types for each tissue (frontal cortex - top panels; hippocampus - bottom panels) across different timepoints (24hr - left panels, 7-day - middle panels, and 21-day - right panels). (b) Dotplots showing the quantification of transcriptomic shifts using a Euclidean distance measure between groups (TBI effect between mTBI and sham groups in red, T4 treatment effect between T4 and mTBI groups in cyan) in individual cell types for each tissue (frontal cortex - left, hippocampus - right) and timepoints (24hr - top panel, 7-day - middle panel, 21-day - bottom panel). The log fold change (logFC) on the y-axis quantifies the global transcriptome shift in the Euclidean distance space between treatment groups, compared to null distribution (randomly shuffling cells between groups and re-calculating Euclidean distance). Each dot is colored by treatment (mTBI vs. sham in red; T4 vs. mTBI in cyan). The size of each dot relates to the adjusted p-value in the Euclidean distance analysis and statistical significance, with colored dots reaching an adjusted p-value < 0.05 whereas gray points not achieving statistical significance. (c) Comparison of DEG counts for mTBI effect (mTBI vs. sham), T4 effect (T4 vs. mTBI), and shared between mTBI and T4 effects, in each cell type in the frontal cortex (left) and hippocampus (right) across timepoints (24hr - top panel, 7-day - middle panel, 21-day - bottom panel).

### The spatiotemporal dynamics of differentially expressed genes (DEGs) affected by mTBI and T4 treatments

To complement the global transcriptome analysis above which captures cumulative, less selective effects on the transcriptome, we identified significant DEGs at false discovery rate (FDR)<0·05 in individual cell types, tissues, and timepoints in response to mTBI (mTBI vs. sham; red bars in Fig.3c; all DEG lists in Supplementary TableS1) or T4 treatment (T4 vs. mTBI; cyan bars in Fig.3c; all DEG lists in Supplementary TableS2). These DEGs represent changes in specific genes with more prominent effect sizes. Glial cells including oligodendrocytes, astrocytes, and microglia consistently had large numbers of DEGs in both the hippocampus and cortex across all timepoints after mTBI (Fig.3c, red bars). The mTBI effects on neurons in the hippocampus were the highest at the early timepoint (24hr) but decreased with time. However, cortical neurons had the largest number of DEGs at 7-day post mTBI. mTBI also induced DEGs in various neuronal subpopulations in both tissues, particularly at the 24hr and 7-day timepoints (Supplementary Fig.S2c; full DEG list in Supplementary TableS3). We note, however, the number of significant DEGs was small in these subtypes of neurons, likely due to the limited sample sizes for each neuronal subtype.

T4 treatment exhibited strong effects on endothelial cells and glial cells (oligodendrocytes, astrocytes, and microglia) in both brain regions as reflected by the higher DEG numbers for these cell types (cyan bars in Fig.2c). T4 treatment also showed differential temporal dynamics in the two tissues evaluated: for the hippocampus the highest number of T4 DEGs were observed at the 7-day stage, particularly in mature oligodendrocytes and OPCs; for the cortex the number of T4 DEGs increased with time, with 21-day having the highest number of DEGs, particularly in mature oligodendrocytes, endothelial cells and microglia (cyan bars in Fig.2c). In neuronal subpopulations, however, few DEGs were affected by T4 treatment (cyan bars in Supplementary Fig.S2c; Supplementary TableS3).

Comparing the DEGs between the mTBI effect and T4 effect, we found that T4 treatment had smaller DEG numbers compared to the mTBI treatment effect (cyan bars compared to red bars in Fig.3c, Supplementary Fig.S2b), but many of the T4 DEGs overlapped with those affected by mTBI for the majority of the cell types (black bars in Fig.3c). These results suggest that T4 treatment targets a subset of genes that were affected by mTBI. We note that differences in the cell numbers were not a major confounder of the DEG numbers in this comparison, as similar numbers of cells were studied from mTBI and T4 groups for each cell type/subtype (Supplementary Fig.S3).

### T4 reversed select DEGs affected by mTBI

There were numerous mTBI-affected genes whose expression levels were reverted closer to the sham level after T4 administration in both the hippocampus (Fig.4a) and the frontal cortex (Fig.4b).

**Fig. 4.**
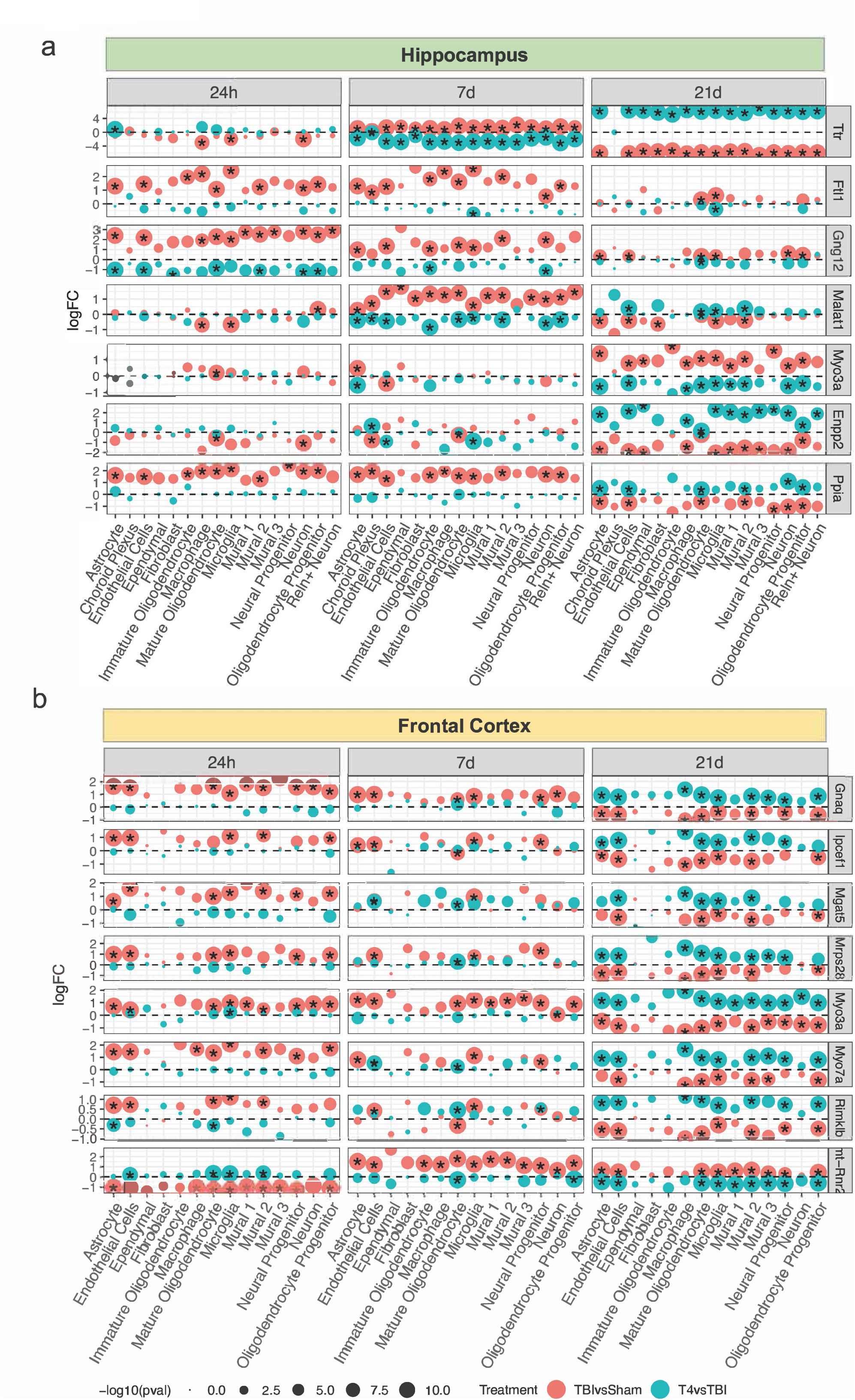
The reversal of mTBI-induced DEGs by T4 treatment across cell types and timepoints in hippocampus and frontal cortex. (a) Select reversed DEGs shared across cell types in the hippocampus. (b) Select reversed DEGs shared across cell types in the frontal cortex. Each row depicts a DEG and each column indicates a cell type, with cell types indicated on the x-axis. The genes which are significantly differentially expressed (adjusted p-value < 0·05) in specific cell types are indicated by a star and the size of the dot corresponds to the −log10 (p-value). The color of each dot indicates TBI effect (red) or T4 effect (cyan). Data for different timepoints are shown in different blocks (24hr - left block; 7-day - middle block, 21-day - right block). The y-axis is the log (fold change) of the gene between groups.

Among the hippocampal DEGs, the T4 transporter transthyretin (*Ttr*) was upregulated by mTBI at the subacute phase across all the cell types of the hippocampus, and was reversed by T4 treatment. Interestingly, at the subchronic phase it was downregulated by mTBI across all cell types except choroid plexus, but upregulated after T4 treatment. Other top mTBI-affected DEGs that were altered across cell types and reversed by T4 included G protein subunit gamma 12 (*Gng12*), metastasis associated lung adenocarcinoma transcript 1 (*Malat1*), myosin IIIA (*Myo3a*) and Peptidylprolyl isomerase A *(Ppia)*. Interestingly, the degree and direction of changes for many of these genes are not necessarily consistent across timepoints. Ferritin light polypeptide 1 (*Ftl1*) was significantly altered by mTBI in the majority of the hippocampal cells at 24hr and 7-day timepoints, but was only reversed by T4 in microglia at 7-day and 21-day timepoints. *Ftl1* plays an important role in iron metabolism and storage, and abnormal iron levels have been shown to be linked to neurodegenerative diseases.^35^ Human FTL, orthologous to mouse *Ftl1*, is upregulated in activated microglia in Alzheimer’s Disease patients.^36^ Notably, *Ftl1* was consistently upregulated by mTBI but downregulated by T4 treatment in microglia across all the timepoints, suggesting that T4 administration restored iron homeostasis. *Gng12* encodes a protein that belongs to the Guanine nucleotide-binding protein (G protein) family and has a role in inflammation and cancer,^37,38^ mTBI increased *Gng12* but T4 suppressed this gene across cell types and timepoints, likely mitigating inflammation. *Malat1* encodes a long non-coding RNA that is highly abundant in the nervous system and plays a role in neuronal cell injury;^39^ this gene was increased by mTBI and decreased by T4 at 7-day but the direction was opposite at 24hr and 21-day post-injury. *Myo3a* is a cytoskeleton gene and variants are related to hearing and vision loss,^40,41^ and this gene was primarily increased by mTBI but inhibited by T4 at 21-day post-injury. *Ppia* mediates inflammation and has shown both beneficial effects intracellularly^42^ and detrimental function extracellularly.^43^ The autoantibody to Ppia is also identified in the serum of mice with TBI^44^ and humans with spinal cord injury.^45^ *Ppia* was increased by mTBI at 24hr and 7-day timepoints but was decreased at 21-day, and T4 increased this gene at 21-day.

In the cortex, top consistent DEGs across cell types and timepoints that were reversed by T4 included *Gnaq, Ipcef1, Mgat5, Mrps28, Myo3a, Myo7a, Rimklb,* and *mt-Rnr2* (Fig.4b). All of these genes except *mt-Rnr2* were induced by mTBI at 24hr in 5-10 cell types, this induction persisted but was weaker at the 7-day timepoint, until these genes were finally downregulated at 21-days. These same genes’ expression patterns were reversed by T4 at 21-days. *mt-Rnr2* exhibited a unique pattern where it was repressed by mTBI at 24hr in 11 cell types, then induced at 7-days in 12 cell types, and induced to a lesser extent at 21-days. T4 reversed the expression of *mt-Rnr2* in 9 cell types at 21-days. These genes encompass a wide variety of functions. *Gnaq* is part of a family of heterotrimeric G protein subunits involved in the negative regulation of neuronal excitability, and when deleted along with *G_11_* in the forebrains of mice, leads to increased seizure susceptibility, neuronal degeneration and reactive gliosis in the hippocampal CA1 region.^46^ *G_q_*/*G_11_* signaling in thyroid follicular cells is also necessary for TSH-induced thyroid hormone synthesis and release.^47^ *Ipcef1*, interaction protein for cytohesin exchange factors 1, is induced in dorsal root ganglion in response to nerve injury.^48^ *Mgat5*, a member of the glycosyltransferase family, is involved in early brain development and differentiation of neural stem and progenitor cells.^49^ *Myo3a* and *Myo7a* are implicated in regulating actin bundles.^50^ As energy needs are high directly post mTBI, we found that *Mrps28* encoding the mitochondrial ribosomal protein 28, and *mt-Rnr2*, mitochondrially encoded 16S RNA were both affected, although in opposite directions. *mt-Rnr2* is also known as Humanin, which suppresses Alzheimer’s disease-related neurotoxicity^51^ and was shown to mitigate mTBI by restoring energy metabolism in our recent study.^9^ The fact that T4 treatment also modulates *mt-Rnr2* supports some shared mechanisms between the two.

The interesting individualized dynamics and tissue/timepoint specificity of each gene suggested that at different injury stages, the decision to enhance or inhibit a gene target requires careful consideration. The largely reversed effect of T4 treatment on these mTBI-affected genes with diverse functions from cell/energy metabolism and iron storage to inflammation, neuronal excitability, and brain development, supports that T4 restores a broad spectrum of functions perturbed by mTBI across cell types and tissues.

### Functional annotation of DEGs affected by mTBI and T4

To further infer the functions of the DEGs play in TBI pathogenesis and T4 intervention, we conducted pathway enrichment analysis using KEGG,^52^ Reactome,^53^ Biocarta, and Gene Ontology^54^ databases. We found diverse pathways related to cellular metabolism, metal ion homeostasis, immune system, inflammation, hypoxia, neurogenesis and neuronal functions to be enriched among the mTBI-affected DEGs, and these pathways showed a dynamic pattern in the two tissues across the three injury stages (Fig.5; Supplementary Fig.S4, TableS4). In the hippocampus, the 24hr acute phase post-TBI exhibited a mixture of upregulated and downregulated pathways; the subacute (7-day) phase showed mainly downregulated pathways; the 21-day subchronic phase saw mostly upregulated pathways (Supplementary Fig.S4). In the cortex, upregulated and downregulated pathways were more evenly distributed in the 24hr acute phase, while downregulated pathways were dominant in the 7-day subacute and 21-day subchronic timepoints (Supplementary Fig.S4).

**Fig. 5.**
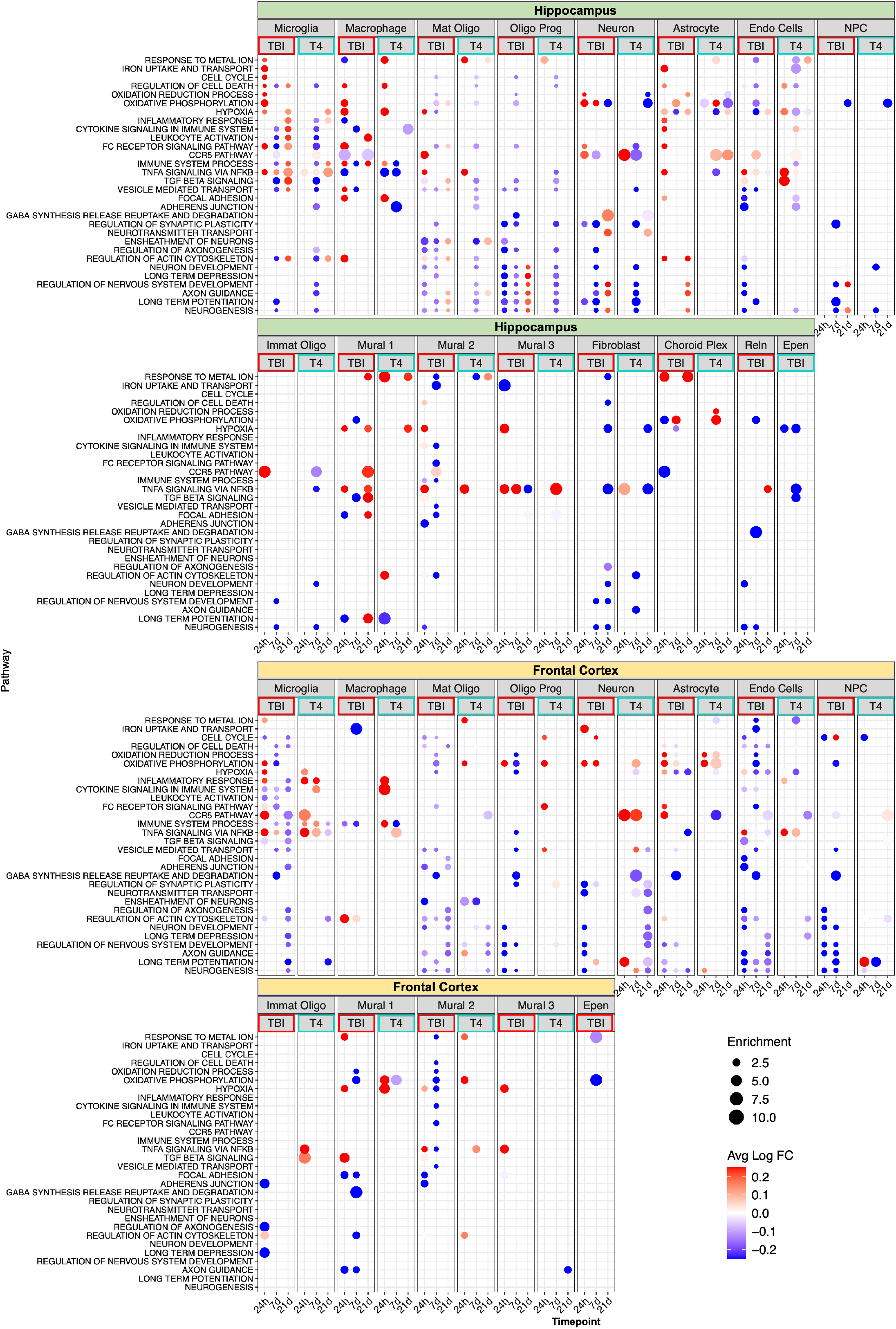
Top enriched pathways responsive to both mTBI or T4 treatments across timepoints and cell types. Each dot is colored by the average fold change between treatment groups (mTBI vs. sham for mTBI effect; T4 vs. TBI for T4 effect) for that cell type for significant DEGs which overlap the indicated pathway. The size of each dot is proportional to the enrichment score. Red indicates upregulation and blue indicate downregulation.

Compared to the mTBI effect on pathways, T4 treatment DEGs had fewer enriched pathways (Supplementary Fig.S5, TableS5). These pathways were related to cell metabolism, cell cycle, glial differentiation, hypoxia, ion homeostasis, cytoskeleton, immune/inflammatory pathways, and neuronal functions. In the hippocampus, T4 upregulated more pathways at 24hr but downregulated most pathways at 7-days and 21-days; in the cortex, T4 treatment upregulated more pathways at 24hr and 7-days, but mostly downregulated pathways at the 21-day stage.

We found numerous metabolic, inflammatory, and neuronal pathways affected by mTBI were also affected by T4 treatment (Fig.5), but the directions of the average expression changes of the DEGs in these pathways were variable depending on the tissue, cell type, and timepoint. For example, nervous system related pathways were downregulated in neural cell types of both brain regions during acute and subacute phases by mTBI. T4 treatment reversed the inhibition of select pathways such as long-term potentiation in the cortical neurons and neural progenitor cells (NPCs) at the acute phases but not in the other cell types or timepoints. Cell metabolism pathways such as oxidative phosphorylation also showed highly spatiotemporal dynamics involving upregualation in certain cell types and timepoints and downregulation in the others by both mTBI and T4 treatment. The upregulation in cortical neurons and astrocytes during acute and subacute phases post-TBI was further enhanced by T4 treatment, but this trend does not necessarily hold true for other cell types. Immune related pathways also had nuanced dynamics. For example, at 24hr post-TBI, immune pathways were extensively elevated across immune cells of both brain regions, and T4 treatment further elevated these pathways in select cell types in the cortex, but less so in the hippocampus. In the subacute (7-day) phase, mTBI suppressed immune pathways in both the cortex and hippocampus post-TBI, whereas T4 treatment increased these pathways in the cortex but further downregulated them in the hippocampus. Some of these pathways, such as oxidative phosphorylation in energy metabolism and long-term potentiation, could be protective responses to mTBI and further enhancement by T4 in select cell populations may help improve the TBI outcome.

### Cell type-specific gene regulatory networks reveal recovery of select gene network programs with T4 treatment

To understand gene regulatory dynamics across treatments and timepoints, we constructed cell type-specific gene regulatory networks (GRNs) with SCING^25^ and detected GRN subnetworks or modules through Leiden clustering^55^ (Supplementary TableS6). Unlike pathway analysis of DEGs which group genes based on known functional categories, GRN analysis offers complementary data-driven information on regulatory relationships among genes and how T4 may counteract mTBI-perturbed gene networks.

Using a linear regression on GRN module expression with treatment and timpoint as independent variables, we identified GRN modules with significantly different expression between mTBI and sham groups but no significant difference between T4-treated and sham mice (Fig.6a). This suggests that T4 intervention in mTBI recovers certain gene programs to a level similar to that of sham mice. These GRN modules were mainly identified from cortical astrocytes and oligodendrocytes as well as from hippocampal oligodendrocytes and oligodendrocyte progenitor cells (OPCs). Pathway analysis on the genes in these GRN modules revealed enrichment for various biological processes. Module5 from hippocampal OPCs was enriched for cell cycle, stress response, and proteasome activity, while Module0 from cortical astrocytes was enriched for vesicle-mediated transport, gene regulation, and RNA splicing (Fig.6a, Supplementary TableS6). As an example, overlaying mTBI v. Sham and T4 v. mTBI DEGs with the hippocampal OPC Module5 showed dynamic upregulation of genes in this subnetwork by mTBI at 24hr (Fig.6b), 7-days (Fig.6c), and 21-days (Fig.6d). T4 treatment mostly downregulated a different subset of genes in this highly connected GRN subnetwork at 7-days post-injury (Fig.6c) and reversed a few select genes at 24hr and 21-day timepoints, likely to counteract dysregulation and restore the balance in network activity.

**Fig. 6.**
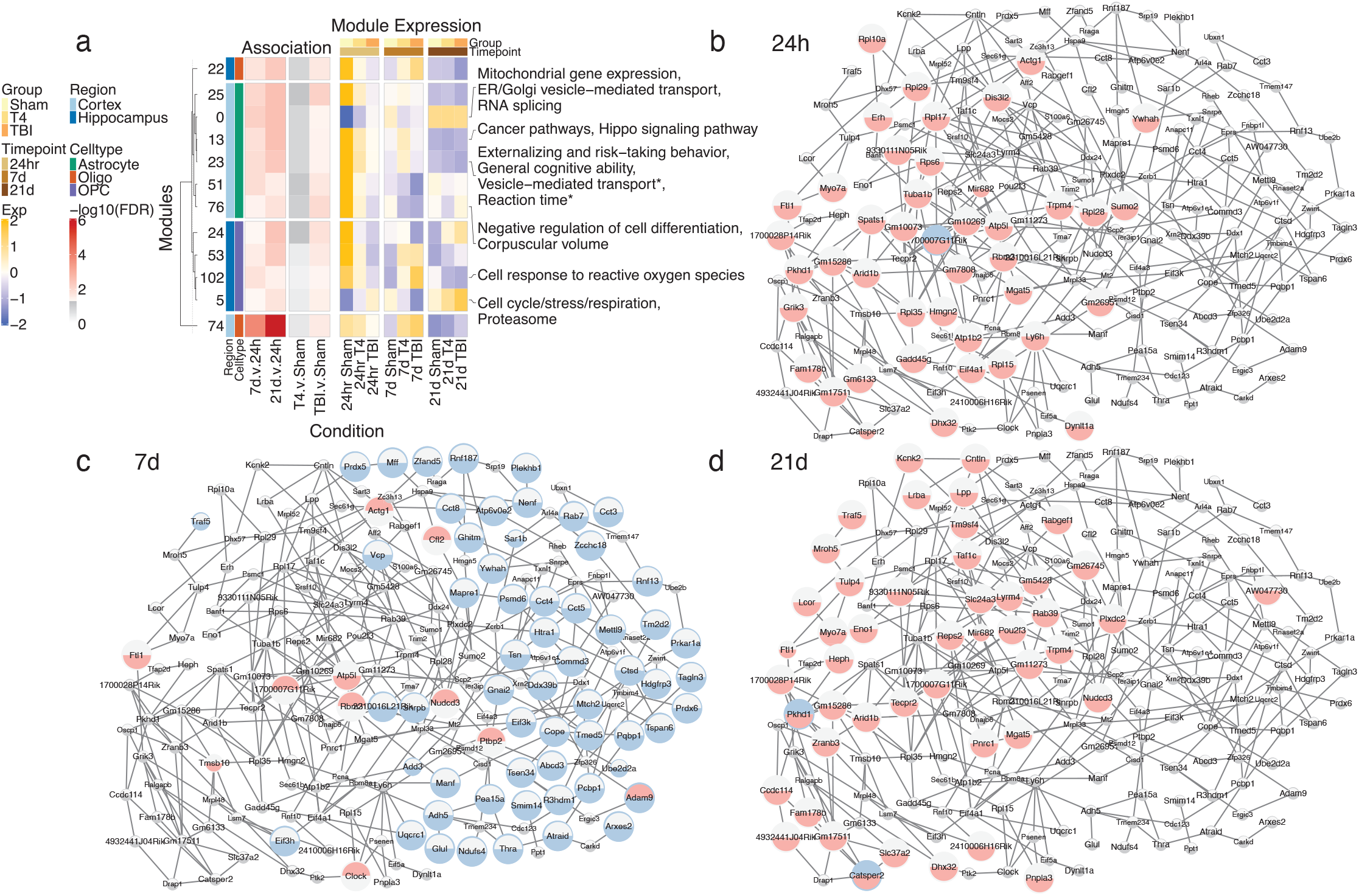
Cell type module association across treatments and timepoints. (a) Heatmap of treatment and time point association to module expression (left), and average module expression for each time point and treatment (right). Asterisks next to pathway or GWAS trait names indicate suggestive enrichment (FDR < 0·1) whereas other pathways listed are at FDR<0·05. (b-d) Hippocampus Oligodendrocyte Precursor cell type module associated with cell cycle and stress response across time points. Larger nodes denote DEGs. Bottom and top halves of network nodes represent DEG direction of mTBI effect (mTBI vs. Sham) and T4 effect (T4 vs. mTBI), respectively. Red, blue, and gray colors represent up, down, and no regulation, respectively.

### Cell-cell communication network analysis identifies key cell type-specific and cross-cell-type shared ligands targeting top DEGs of mTBI and T4 treatment effects

To characterize the interactions between different cell types impacted by mTBI and/or T4 treatment, we identified ligands that were predicted to target large numbers of significant DEGs in key cell types affected by mTBI and/or T4, including mature oligodendrocytes, astrocytes, endothelial cells, NPCs, and neurons in hippocampal and cortical tissue (Fig.7). In both the hippocampus and cortex, ligands shared across cell types that were predicted for DEGs from each comparison (TBI vs. sham and T4 vs. TBI) and for every time point (24hr, 7-days, 21-days) included β-Amyloid Precursor Protein (App), High-mobility Group Box-1 (HMGB1), Ribosomal Protein S19 (Rps19), and Pleiotrophin (Ptn) (Fig.7a-d for hippocampus; 7e-h for frontal cortex). Each of these ligands were predicted to target DEGs shared between mTBI effect (TBI vs. sham) and T4 treatment effect (T4 vs. TBI), but the patterns varied between ligands. For example, App was more strongly downregulated by mTBI across cell types and timepoints in the hippocampus (Fig.7a) with a weaker trend in the cortex (Fig.7e), and both tissues displayed a mixture of up- or downregulation by T4 that varied between timepoints (Fig.7a, 7e). Ligands Hmgb1, Rps19, and Ptn demonstrated a different trend, with their expression levels mostly upregulated by mTBI during the 24hr acute and 7-day subacute phases in both brain regions but trended more towards downregulation by T4 across timepoints. Visualization of the cross-cell-type interactions mediated by these ligands (Fig.7b-d for hippocampus; Fig.7f-h for frontal cortex) also support that they participate in regulating not only the shared DEGs between mTBI and T4 effects (marked by yellow tracks), but also mTBI- (marked by red tracks) or T4-specific DEGs (marked by blue tracks). Aggregation of β-amyloid, the product of App cleavage, is a one of the pathological hallmarks of Alzheimer’s disease, and our cell-cell communication analyses revealed App in relation to genes impacted by mTBI and T4, highlighting the known proposal that TBI is a risk factor for Alzheimer’s disease.^56^ App has been shown to exert neuroprotective effects following TBI by binding to heparan sulfate proteoglycans.^57^ Hmgb1 promotes neuroinflammation and has been linked to the pathogenesis of TBI.^58,59^ Additionally, our data confirmed previous findings that Ptn, a neurotrophic factor critical in neuroregeneration, is upregulated specifically in the subacute phase of TBI.^60^

**Fig. 7.**
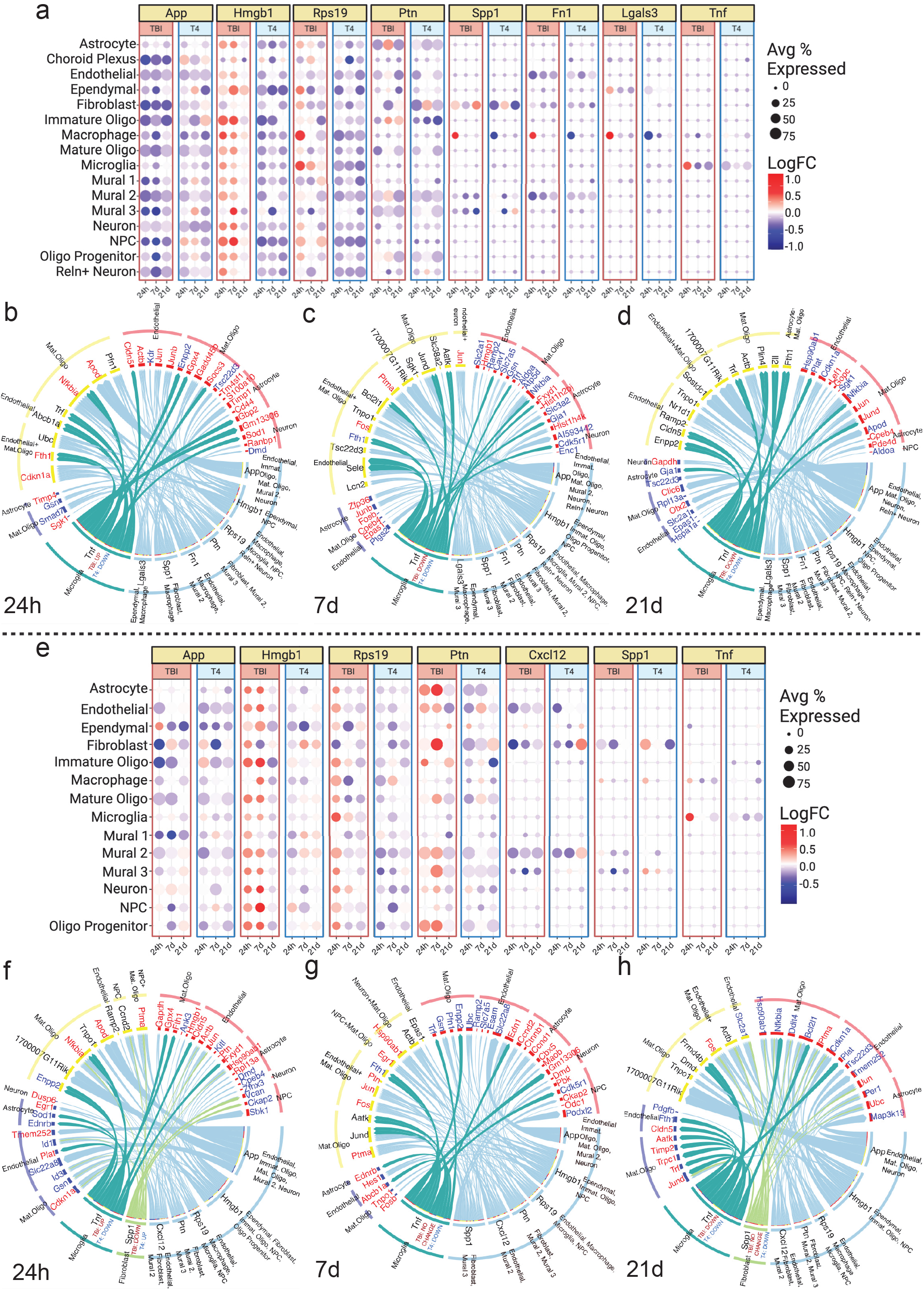
Cell-cell communication between hippocampal and cortical cell types. (a, e) Expression patterns of ligand genes predicted to target mature oligodendrocyte, astrocyte, endothelial, NPC, and neuron DEGs in the hippocampus (a) and frontal cortex (e) for mTBI effect (mTBI vs. sham) and T4 effect (T4 vs. mTBI) at 24 hours, 7-day, and 21-day. Dot colors indicate the log2 fold change (Log2FC) of each ligand for each treatment effect at each timepoint. Dot sizes indicate the percentage of cells expressing the ligands at each condition. (b-d, f-h). Ligands and their corresponding target genes (DEGs) that they are predicted to influence at 24 hours, 7-day, and 21-day in the hippocampus (b-d) and cortex (f-h). The corresponding cell types and target genes for each ligand are listed on the outer border and inner border tracks, respectively. Target genes along the blue track are DEGs from the mTBI effect (mTBI vs. sham) only, genes along the red track are DEGs from the T4 effect (T4 vs. mTBI) only, and genes along the yellow track are shared DEGs between mTBI effect (TBI vs. sham) and T4 effect (T4 vs. mTBI). DEGs are colored based on the direction of the fold-change in the corresponding comparison (red=up, blue=down, black=opposite direction in mTBI effect vs. T4 effect). The expression change for cell type-specific ligands is listed for the mTBI effect (mTBI vs. sham) and T4 effect (T4 vs. mTBI) below the ligand name. Ligands from multiple cell types are not labeled with the direction of change information as the directions can be different between cell types.

Other more cell-type specific ligands such as Osteopontin (Spp1) and Tumor Necrosis Factor (Tnf) were also found to be shared between brain regions and treatment effects. Spp1 was predicted to be a ligand sent by fibroblasts, macrophages, and mural cells in both brain regions, but the direction of changes in Spp1 varied between cell types and tissues, presenting a more nuanced fine-tuning of Spp1 modulated cell-cell communications depending on the cell type and tissue context. Nevertheless, mTBI and T4 induced opposite directions of change in Spp1 expression, supporting that T4 treatment opposed the mTBI-induced changes. At all timepoints in the hippocampus and cortex, Tnf was a microglia-specific ligand targeting primarily shared DEGs between mTBI effect and T4 treatment effect (genes listed in yellow track), as well as DEGs with mTBI-specific effect (genes in red track) in the hippocampus (Fig.7b-d). In the frontal cortex, however, Tnf was predicted to additionally target DEGs with T4-specific effects (genes in blue track in Fig.7f-h) in various cortical cell types. Proinflammatory factors Spp1 and Tnf were previously identified as TBI-associated hub genes in a TBI rat study, and both genes were upregulated when in the cerebral cortex of the TBI rat model compared to control at four hours post-TBI.^61^

In addition to the above ligands shared between brain regions, we also found tissue-specific ligands including Fn1 and Lgals3 in the hippocampus and Cxcl12 in the cortex, suggesting their tissue-specific regulatory roles. Fibronectin 1 (Fn1) and Galectin 3 (Lgals3) were upregulated by mTBI but downregulated by T4 treatment at the 24hr acute phase in hippocampal macrophages. Fn1 is involved in extracellular matrix and cell adhesion and was also identified as a TBI hub gene and potential TBI biomarker.^61^ Lgals3 is a driver of inflammation and has been shown to be upregulated on the protein level after TBI, but this upregulation was reversed by T4 treatment.^62^ CXC Motif Chemokine Ligand 12 (Cxcl12) was downregulated by mTBI across all timepoints but was upregulated by T4 at the subchronic 21-day timepoint in the fibroblast and mural2 cell populations. Cxcl12 is a key chemokine that can mediate peripheral blood cell recruitment to the TBI brain.^63^ Overall, these various ligands identified may play central roles in orchestrating cell-cell communications for both the mTBI effect and the T4 treatment effect.

### Disease and trait association analysis of genes affected by mTBI and T4 treatment

Lastly, to assess the human pathophysiology relevance of the DEGs identified from mTBI effect (mTBI vs. sham) and T4 effect (T4 vs. mTBI), we evaluated whether they were enriched for candidate genes implicated in human diseases and traits based on the human GWAS catalog.^30^ In both the hippocampus and cortex, many of the cell-type specific DEGs affected by mTBI and/or T4 were enriched for genes involved in brain related diseases and traits such as cognitive performance, depression, insomnia, schizophrenia and Alzheimer’s disease (Fig.8, full list in Supplementary TableS7 and S8), supporting that both the injury itself and the T4 intervention modulate disease-associated genes across cell types. However, key differences between the two treatments were also present. In the hippocampus, oligodendrocyte progenitors and mature oligodendrocytes were among the cell types whose DEGs affected by mTBI across timepoints showed the most enrichment for many brain related diseases and traits. In contrast, T4 treatment affected DEGs were enriched for disease associated genes primarily at the 7-day time point. For the oligodendrocyte progenitors from the cortex, disease gene enrichment was mainly observed at the 24hr time point after mTBI but at 21-days after T4 treatment. mTBI DEGs from astrocytes from both the hippocampus and cortex showed enrichment for several diseases, but T4 DEGs had minimal enrichment. In neuronal cells, mTBI DEGs from both tissues showed significant enrichment of several brain related disorders. In contrast, significant disease gene enrichment was mainly observed for T4 DEGs in cortical neurons but less so for hippocampal neurons. These results suggest both shared and distinct effects of mTBI and T4 treatments on disease phenotypes.

**Fig. 8.**
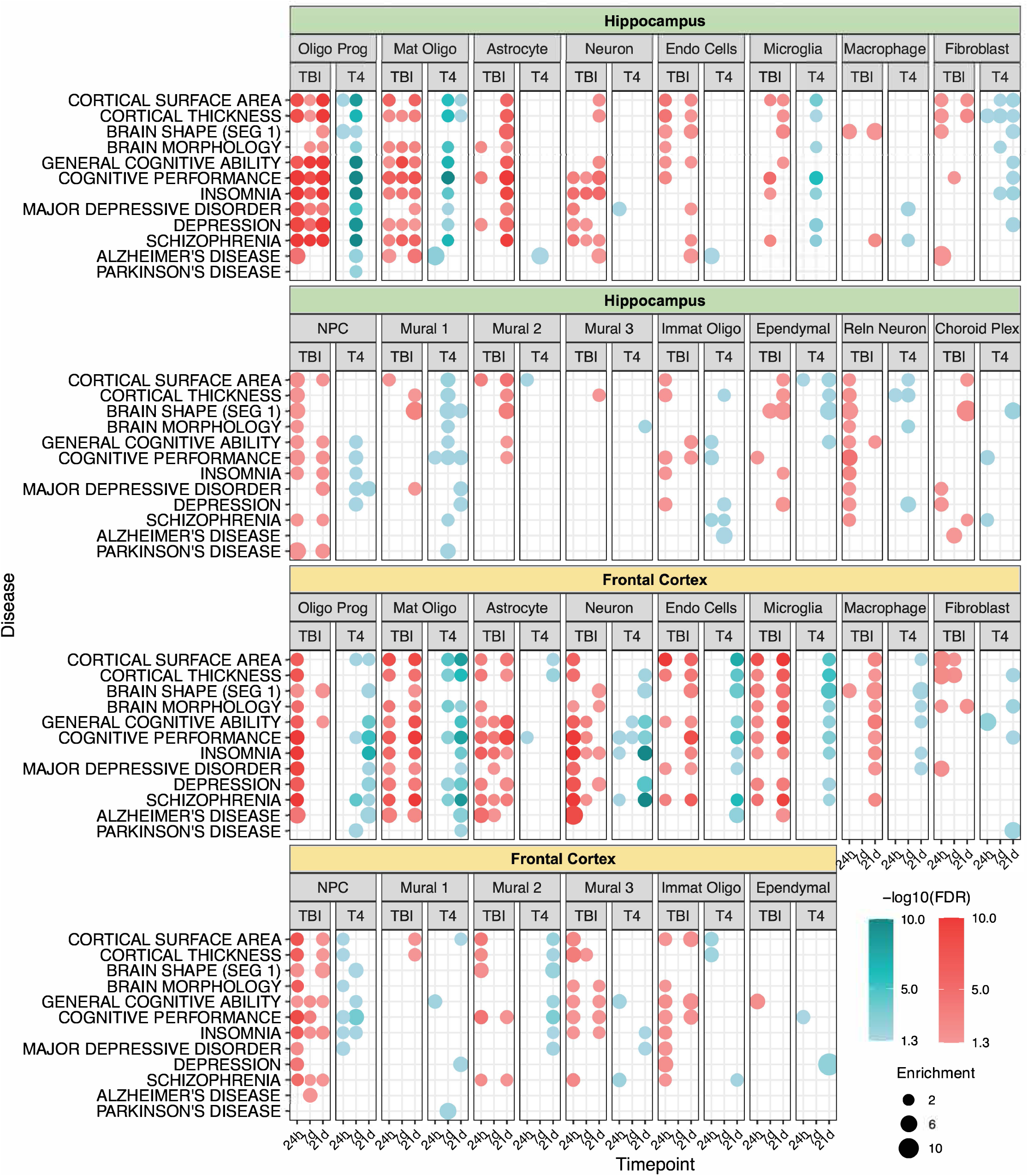
Human disease/trait enrichment analysis of genes responsive to mTBI and T4 across timepoints and tissues presented in a dot plot. Brain disease related candidate gene sets were retrieved from the GWAS catalog, which were used for enrichment analysis of the DEGs affected by mTBI or T4 treatment (p-value <0·01). A disease/trait was considered to be significantly enriched for mTBI/T4 DEGs using FDR <0·05 and gene overlap number >= 3 as cutoffs. The size of the dots is proportional to the fold enrichment score. For visualization purposes the enrichment limit was set to 10. Red and blue colors were used to display the disease enrichment results for mTBI DEGs and T4 DEGs, respectively. Color shades correspond to the −log10(FDR). Cell type abbreviations: Oligodendrocyte progenitor (Oligo Prog), Mature Oligodendrocyte (Mat Oligo), Endothelial Cells (Endo Cells), Neural Progenitor Cells (NPC), Immature Oligodendrocyte (Immat Oligo).

## Discussion

Due to the complexity of mTBI, effective therapeutics is currently lacking. The thyroid hormone pathway has emerged as a new therapeutic avenue based on recent evidence, including results from our spatiotemporal scRNAseq studies of mTBI.^8,9^ Previously, we showed the ability of acute T4 administration within 6 hours post-mTBI to prevent the cognitive impairment induced by mTBI^8^ and here we replicated this finding in an independent set of experiments. However, how targeting the thyroid hormone pathway mitigates mTBI remains unclear. In this study, using a systems-level approach we investigated the spatiotemporal molecular mechanisms of T4 treatment in two important brain regions (hippocampus and frontal cortex) and three post injury stages (acute, subacute, and subchronic). The current study aims to elucidate which timepoint (acute - 24hr vs. subacute −7-day vs. subchronic - 21-day), brain region (hippocampus vs. frontal cortex), cell types (neuronal or various glial cells), and genes, pathways, and networks are important for the observed beneficial effect of T4 treatment on cognitive recovery.

In terms of temporal sensitivity, we found that while mTBI induced more dramatic effects at the acute and subacute stages, T4 treatment had minimal effect in the acute phase but exhibited more pronounced influence at later stages. This suggests that either the strong effect of mTBI in the acute phase overwhelms the acute effects of T4, or T4 takes time to execute its effect.

Between the two brain regions, we found that T4 treatment induced earlier changes in the hippocampal cell populations starting from the 7-day subacute phase, whereas the changes in the frontal cortex tended to occur later at the 21-day subchronic stage. This is particularly apparent in the cell proportion (Fig.2), global transcriptome shift (Fig.3b), and DEG number (Fig.3c) analyses, where T4 administration induced more changes in the hippocampal cell populations earlier than in the frontal cortex cell types. These results suggest that the recovery of hippocampal functions is faster when T4 is administered, which could be caused by faster and/or more preferred distribution of T4 from the intraperitoneal injection site to the hippocampal region and requires further T4 tracing studies to explain the differences in spatial specificity and dynamics. Notably, *Ttr*, the T4 transporter, was uniquely altered by T4 in the hippocampus but not in the frontal cortex, which may indicate a T4 transport advantage of the hippocampus. Alternatively, hippocampal cell populations may have higher sensitivity to T4 if dosage distribution bias can be excluded.

In terms of cell type selectivity of T4 effect, different analyses offered complementary views. In the cell proportion analysis (Fig.2), T4 primarily counteracted the decrease in Mural3 cells at 7-days and the increase in endothelial cells at 21-days caused by mTBI to normalize these cell types to the sham control levels. More interestingly, T4 uniquely caused an increase in endothelial cells and oligodendrocyte progenitor cells at 7-days, which may enhance BBB repair and replace damaged or dysfunctional mature oligodendrocytes. At 21-days, T4 also uniquely boosted the choroid plexus population, which plays a critical role in neural repair.^64^ These cells are not part of the hippocampus but are typically included during hippocampal dissection.^33^ All these changes occurred in or near the hippocampus, again supporting that this brain region and its related cell types are main T4 targets. To supplement the cell proportion analysis, we used a Euclidean distance-based method to measure global transcriptomic changes in individual cell types (Fig.3b). Such analysis identifies cell types which may not show cell proportion changes but have major reprogramming in the transcriptome. This analysis revealed significant changes in hippocampal cell types including endothelial cells at 7-day, mature oligodendrocytes at 7- and 21-day, astrocytes at 21-day, as well as cortical cell types including mature oligodendrocytes at 7-day and neurons and neural progenitor cells (NPCs) at 21-day. Lastly, we also assessed cell type sensitivity based on the significant DEG counts from individual cell types (Fig.3c). This analysis mostly identifies genes with more prominent effect sizes and showed that the largest numbers of DEGs were from mature oligodendrocytes, endothelial cells, and microglia from both brain regions and cross timepoints. Taken together, these various complementary methods consistently identified mature oligodendrocytes and endothelial cells as sensitive cell types to T4 treatment. Additional T4 target cells identified from at least one of the three methods include OPCs, mural cells, choroid plexus, and astrocytes in or near the hippocampus, microglia from both brain regions, and cortical neuronal populations. Many of these cell types are important for injury and neural repair. For instance, oligodendrocytes form the myelin assembly in the CNS, wrapping around neuron axons, which ensures the fast conduction of nerve impulse^65^ and metabolic support.^66,67^ TBI could cause the progressive demyelination and degeneration of axons.^68^ Remyelination is fulfilled mainly by newly generated oligodendrocytes,^69^ and OPCs contribute to remyelination and glial scar formation.

To address which genes, pathways, and networks are modulated by T4 treatment and which ones may be responsible for the protective effect of T4, we also carried out multiple complementary analyses. First, based on DEGs identified from individual cell types at each timepoint, we found interesting genes whose expression was altered by mTBI but the change was reversed by T4 across cell types and timepoints. These genes were found to be involved in diverse functions including iron metabolism/storage (*Ftl1*), cell/energy metabolism and mitochondria function (*Ttr*, *mt-Rnr2, Mrps28*), inflammation (*Ggn12, Ppia*), neuronal activity (*Gnag*), neural differentiation (*Mgat5, Malat1*), and cytoskeleton (*Myo3a, Myo7a*, *Ipcef1*). Similar functional categories were also captured in the pathway enrichment analysis of the mTBI-affected DEGs that were also modulated by T4 treatment. We also supplemented these standard analyses with advanced network modeling approaches, including both within-cell-type GRNs and cell-cell communication networks, to identify network targets and regulators. GRN analysis showed that specific subnetworks from cortical astrocytes and oligodendrocytes as well as from hippocampal oligodendrocytes and OPCs were upregulated by mTBI but were normalized to control levels by T4 treatment. As cell-type GRN is not based on DEGs but gene regulatory relationships within each cell type, this analysis revealed additional processes compared to the DEG and pathway analyses, such as stress response, cell cycle, vesicle transport, transcriptional regulation within the select glial cells to be mitigated by T4 treatment. Lastly, our cell-cell communication analysis revealed potential secreted ligands from numerous cell types that likely regulate cross-cell-type gene regulation that is important for both mTBI and T4 actions. These ligands include numerous inflammation regulators Hmgb1, Spp1, Tnf, Lgal3, and Cxcl12, important neuroplasticity factors App and Ptn, and a glycoprotein of the extracellular matrix and adhesion molecule Fn1. T4 treatment reversed the expression levels of several of these ligands, suggesting that T4 may normalize the TBI-perturbed cell-cell communications by modulating these regulatory ligands. These diverse gene, pathway, and network level analyses provide insight into the broad molecular functions of T4 in the brain which likely underlie the efficacy of T4 treatment.

Our studies were carried out in a mouse model of mTBI. To assess the relevance of our results to human pathophysiology, we also integrated our mTBI and T4 DEGs with candidate disease genes implicated in human GWAS. This analysis confirmed that the DEGs affected by the injury itself and those modulated by T4 treatment are relevant to cognitive performance, psychiatric disorders such as depression and schizophrenia, and neurodegenerative disorders such as Alzheimer’s disease. These results suggest that T4 treatment may be able to alleviate these clinical conditions known to be associated with TBI due to the significant gene overlaps between our T4 mouse study and human diseases.

In summary, our investigation across tissues and timepoints offers a comprehensive spatiotemporal understanding of the cellular and molecular mechanisms in T4 induced reversal of mTBI dysfunction using single cell technologies. Our data showed that glial cells such as oligodendrocytes, astrocytes, microglia, OPCs, endothelial cells, and mural cells as well as choroid plexus near hippocampus and neuronal populations in the frontal cortex to be particularly important for T4-dependent treatment effects, many of which were among the vulnerable cell types in mTBI. Diverse genes and pathways related to cell metabolism, iron homeostasis, immune response, cytoskeleton, and nervous system were affected by T4 treatment, and these pathways showed brain-region, cell-type, and injury stage specificity and dynamics. Comparison between the T4 effect and mTBI effect on individual cell types in both brain regions revealed the convergence of numerous DEGs, pathways, gene subnetworks, cell crosstalk, and disease/trait association that likely underlie the protective effect of T4 treatment on cognitive and neuropsychiatric functions after mTBI. Our results provide molecular support that T4 administration affects a broad spectrum of genes, biological processes, and networks to prevent the progression of mTBI-induced brain dysfunction and diseases. This broader effect of T4 may offer an improved efficacy over other therapeutic options that focus on specific pathways and targets. We acknowledge that future studies are warranted to further understand which of the cell types, pathways, and regulators play a causal role in the protective effect of T4 and which ones may cause unwanted side effects.

## Contributors

X.Y. and F.G-P. conceived the idea. G.Z., G.D., J.C., M.C., S.S.W., and X.Y. drafted the manuscript. G.Z., G.D., J.C., M.C., S.S.W., K.D.A., and X.Y. reviewed and edited manuscript, which was approved by all authors. G.Z., V.P.S., and Z.Y. conducted animal experiments. G.D., G.Z., I.S.A. and I.C. performed scRNA-seq. G.Z., J.C., M.C., D.A, G.D., N.P., and N.W. directly accessed and verified the data reported in the manuscript, and performed bioinformatic analyses of scRNA-seq data. G.Z., G.D., J.C., M.C., and K.A. generated figures.

## Acknowledgements

X.Y. and F.G-P. are funded by R01 NS117148. We thank Daniel Sung Min Ha for uploading the sequencing data to GEO.

## Data Sharing Statement

Raw RNA-seq data used in this study have been deposited in the NCBI Gene Expression Omnibus (GEO) data sets.

## References

1 Dewan MC, Rattani A, Gupta S, Baticulon RE, Hung YC, Punchak M, et al. Estimating the global incidence of traumatic brain injury. J Neurosurg 2018; 130. DOI:10.3171/2017.10.JNS17352.

2 Friede A, Reid JA, Ory HW. CDC WONDER: a comprehensive on-line public health information system of the Centers for Disease Control and Prevention. Am J Public Health 1993; 83: 1289.

3 Dixon KJ. Pathophysiology of Traumatic Brain Injury. Phys Med Rehabil Clin N Am 2017; 28: 215– 25.

4 Jamora CW, Young A, Ruff RM. Comparison of subjective cognitive complaints with neuropsychological tests in individuals with mild vs more severe traumatic brain injuries. Brain Inj 2012; 26: 36–47.

5 Wallace EJ, Mathias JL, Ward L. Diffusion tensor imaging changes following mild, moderate and severe adult traumatic brain injury: a meta-analysis. Brain Imaging Behav 2018; 12: 1607–21.

6 Dams-O’Connor K, Juengst SB, Bogner J, Chiaravalloti ND, Corrigan JD, Giacino JT, et al. Traumatic brain injury as a chronic disease: insights from the United States Traumatic Brain Injury Model Systems Research Program. Lancet Neurol 2023; 22: 517–28.

7 Izzy S, Chen PM, Tahir Z, Grashow R, Radmanesh F, Cote DJ, et al. Association of Traumatic Brain Injury With the Risk of Developing Chronic Cardiovascular, Endocrine, Neurological, and Psychiatric Disorders. JAMA Netw Open 2022; 5: e229478.

8 Arneson D, Zhang G, Ying Z, Zhuang Y, Byun HR, Ahn IS, et al. Single cell molecular alterations reveal target cells and pathways of concussive brain injury. Nat Commun 2018; 9: 3894.

9 Arneson D, Zhang G, Ahn IS, Ying Z, Diamante G, Cely I, et al. Systems spatiotemporal dynamics of traumatic brain injury at single-cell resolution reveals humanin as a therapeutic target. Cell Mol Life Sci 2022; 79: 480.

10 Hiebert JB, Shen Q, Thimmesch AR, Pierce JD. Traumatic brain injury and mitochondrial dysfunction. Am J Med Sci 2015; 350: 132–8.

11 Doğanyiğit Z, Erbakan K, Akyuz E, Polat AK, Arulsamy A, Shaikh MF. The Role of Neuroinflammatory Mediators in the Pathogenesis of Traumatic Brain Injury: A Narrative Review. ACS Chem Neurosci 2022; 13: 1835–48.

12 Ng SY, Lee AYW. Traumatic Brain Injuries: Pathophysiology and Potential Therapeutic Targets. Front Cell Neurosci 2019; 13: 528.

13 Ray SK, Dixon CE, Banik NL. Molecular mechanisms in the pathogenesis of traumatic brain injury. Histol Histopathol 2002; 17: 1137–52.

14 Bunevicius A. Chapter 25 - Thyroid hormone actions in traumatic brain injury. In: Rajendram R, Preedy VR, Martin CR, eds. Cellular, Molecular, Physiological, and Behavioral Aspects of Traumatic Brain Injury. Academic Press, 2022: 305–16.

15 Crupi R, Paterniti I, Campolo M, Di Paola R, Cuzzocrea S, Esposito E. Exogenous T3 administration provides neuroprotection in a murine model of traumatic brain injury. Pharmacol Res 2013; 70. DOI:10.1016/j.phrs.2012.12.009.

16 Genovese T, Impellizzeri D, Ahmad A, Cornelius C, Campolo M, Cuzzocrea S, et al. Post-ischaemic thyroid hormone treatment in a rat model of acute stroke. Brain Res 2013; 1513: 92–102.

17 Brewer GJ, Torricelli JR. Isolation and culture of adult neurons and neurospheres. Nat Protoc 2007;2: 1490–8.

18 Macosko EZ, Basu A, Satija R, Nemesh J, Shekhar K, Goldman M, et al. Highly parallel genome-wide expression profiling of individual cells using nanoliter droplets. Cell 2015; 161: 1202–14.

19 Hao Y, Hao S, Andersen-Nissen E, Mauck WM 3rd, Zheng S, Butler A, et al. Integrated analysis of multimodal single-cell data. Cell 2021; 184: 3573–3587.e29.

20 Blondel VD, Guillaume JL, Lambiotte R, Lefebvre E. Fast unfolding of communities in large networks. Journal of Statistical Mechanics-Theory and Experiment 2008.

21 Han X, Wang R, Zhou Y, Fei L, Sun H, Lai S, et al. Mapping the Mouse Cell Atlas by Microwell-Seq. Cell 2018; 172: 1091–1107 e17.

22 Zeisel A, Muñoz-Manchado AB, Codeluppi S, Lönnerberg P, La Manno G, Juréus A, et al. Brain structure. Cell types in the mouse cortex and hippocampus revealed by single-cell RNA-seq. Science 2015; 347: 1138–42.

23 Bhattacherjee A, Djekidel MN, Chen R, Chen W, Tuesta LM, Zhang Y. Cell type-specific transcriptional programs in mouse prefrontal cortex during adolescence and addiction. Nat Commun 2019; 10: 4169.

24 Saunders A, Macosko EZ, Wysoker A, Goldman M, Krienen FM, de Rivera H, et al. Molecular Diversity and Specializations among the Cells of the Adult Mouse Brain. Cell 2018; 174: 1015–1030.e16.

25 Littman R, Cheng M, Wang N, Peng C, Yang X. SCING: Inference of robust, interpretable gene regulatory networks from single cell and spatial transcriptomics. iScience 2023; 26: 107124.

26 Aibar S, González-Blas CB, Moerman T, Huynh-Thu VA, Imrichova H, Hulselmans G, et al. SCENIC: single-cell regulatory network inference and clustering. Nat Methods 2017; 14: 1083–6.

27 Kuleshov MV, Jones MR, Rouillard AD, Fernandez NF, Duan Q, Wang Z, et al. Enrichr: a comprehensive gene set enrichment analysis web server 2016 update. Nucleic Acids Res 2016; 44: W90–7.

28 Shannon P, Markiel A, Ozier O, Baliga NS, Wang JT, Ramage D, et al. Cytoscape: a software environment for integrated models of biomolecular interaction networks. Genome Res 2003; 13: 2498–504.

29 Browaeys R, Saelens W, Saeys Y. NicheNet: modeling intercellular communication by linking ligands to target genes. Nat Methods 2020; 17: 159–62.

30 Sollis E, Mosaku A, Abid A, Buniello A, Cerezo M, Gil L, et al. The NHGRI-EBI GWAS Catalog: knowledgebase and deposition resource. Nucleic Acids Res 2023; 51: D977–85.

31 Pitts MW. Barnes Maze Procedure for Spatial Learning and Memory in Mice. Bio Protoc 2018; 8. DOI:10.21769/bioprotoc.2744.

32 Peris LR, Scheuber MI, Shan H, Braun M, Schwab ME. Barnes maze test for spatial memory: A new, sensitive scoring system for mouse search strategies. Behav Brain Res 2024; 458: 114730.

33 Olney KC, Todd KT, Pallegar PN, Jensen TD, Cadiz MP, Gibson KA, et al. Widespread choroid plexus contamination in sampling and profiling of brain tissue. Mol Psychiatry 2022; 27: 1839–47.

34 Liu R, Zhang Z, Chen Y, Liao J, Wang Y, Liu J, et al. Choroid plexus epithelium and its role in neurological diseases. Front Mol Neurosci 2022; 15: 949231.

35 Muhoberac BB, Vidal R. Abnormal iron homeostasis and neurodegeneration. Front Aging Neurosci 2013; 5: 32.

36 Kenkhuis B, Somarakis A, de Haan L, Dzyubachyk O, IJsselsteijn ME, de Miranda NFCC, et al. Iron loading is a prominent feature of activated microglia in Alzheimer’s disease patients. Acta Neuropathol Commun 2021; 9: 27.

37 Larson KC, Lipko M, Dabrowski M, Draper MP. Gng12 is a novel negative regulator of LPS-induced inflammation in the microglial cell line BV-2. Inflamm Res 2010; 59: 15–22.

38 Liu R, Liu Z, Zhao Y, Cheng X, Liu B, Wang Y, et al. GNG12 as A Novel Molecular Marker for the Diagnosis and Treatment of Glioma. Front Oncol 2022; 12: 726556.

39 Ma P, Li Y, Zhang W, Fang F, Sun J, Liu M, et al. Long Non-coding RNA MALAT1 Inhibits Neuron Apoptosis and Neuroinflammation While Stimulates Neurite Outgrowth and Its Correlation With MiR-125b Mediates PTGS2, CDK5 and FOXQ1 in Alzheimer’s Disease. Curr Alzheimer Res 2019; 16: 596–612.

40 Gunther LK, Cirilo JA Jr, Desetty R, Yengo CM. Deafness mutation in the MYO3A motor domain impairs actin protrusion elongation mechanism. Mol Biol Cell 2022; 33: ar5.

41 Bueno AS, Nunes K, Dias AMM, Alves LU, Mendes BCA, Sampaio-Silva J, et al. Frequency and origin of the c.2090T>G p.(Leu697Trp) MYO3A variant associated with autosomal dominant hearing loss. Eur J Hum Genet 2022; 30: 13–21.

42 Lee SP, Hwang YS, Kim YJ, Kwon KS, Kim HJ, Kim K, et al. Cyclophilin a binds to peroxiredoxins and activates its peroxidase activity. J Biol Chem 2001; 276: 29826–32.

43 Pasetto L, Pozzi S, Castelnovo M, Basso M, Estevez AG, Fumagalli S, et al. Targeting Extracellular Cyclophilin A Reduces Neuroinflammation and Extends Survival in a Mouse Model of Amyotrophic Lateral Sclerosis. J Neurosci 2017; 37: 1413–27.

44 Harper MM, Rudd D, Meyer KJ, Kanthasamy AG, Anantharam V, Pieper AA, et al. Identification of chronic brain protein changes and protein targets of serum auto-antibodies after blast-mediated traumatic brain injury. Heliyon 2020; 6: e03374.

45 Arevalo-Martin A, Grassner L, Garcia-Ovejero D, Paniagua-Torija B, Barroso-Garcia G, Arandilla AG, et al. Elevated Autoantibodies in Subacute Human Spinal Cord Injury Are Naturally Occurring Antibodies. Front Immunol 2018; 9: 2365.

46 Wettschureck N, van der Stelt M, Tsubokawa H, Krestel H, Moers A, Petrosino S, et al. Forebrain-specific inactivation of Gq/G11 family G proteins results in age-dependent epilepsy and impaired endocannabinoid formation. Mol Cell Biol 2006; 26: 5888–94.

47 Kero J, Ahmed K, Wettschureck N, Tunaru S, Wintermantel T, Greiner E, et al. Thyrocyte-specific Gq/G11 deficiency impairs thyroid function and prevents goiter development. J Clin Invest 2007; 117: 2399.

48 Guan X, Zhu X, Tao Y-X. Peripheral nerve injury up-regulates expression of interactor protein for cytohesin exchange factor 1 (IPCEF1) mRNA in rat dorsal root ganglion. Naunyn Schmiedebergs Arch Pharmacol 2009; 380: 459–63.

49 Yale AR, Kim E, Gutierrez B, Nicole Hanamoto J, Lav NS, Nourse JL, et al. Regulation of neural stem cell differentiation and brain development by MGAT5-mediated N-glycosylation. Stem Cell Reports 2023; 18: 1340.

50 Suter DM. Functions of Myosin Motor Proteins in the Nervous System. Neurobiology of Actin 2011;: 45–72.

51 Tajima H, Niikura T, Hashimoto Y, Ito Y, Kita Y, Terashita K, et al. Evidence for in vivo production of Humanin peptide, a neuroprotective factor against Alzheimer’s disease-related insults. Neurosci Lett 2002; 324: 227–31.

52 Kanehisa M, Goto S. KEGG: kyoto encyclopedia of genes and genomes. Nucleic Acids Res 2000;28: 27–30.

53 Croft D, Mundo AF, Haw R, Milacic M, Weiser J, Wu G, et al. The Reactome pathway knowledgebase. Nucleic Acids Res 2014; 42: D472–7.

54 Gene Ontology, Consortium. Gene Ontology Consortium: going forward. Nucleic Acids Res 2015;43: D1049–56.

55 Traag VA, Waltman L, van Eck NJ. From Louvain to Leiden: guaranteeing well-connected communities. Sci Rep 2019; 9: 5233.

56 Sivanandam TM, Thakur MK. Traumatic brain injury: A risk factor for Alzheimer’s disease. Neurosci Biobehav Rev 2012; 36: 1376–81.

57 Plummer S, den Heuvel C V, Thornton E, Corrigan F, Cappai R. The Neuroprotective Properties of the Amyloid Precursor Protein Following Traumatic Brain Injury. Aging Dis 2016; 7. DOI:10.14336/AD.2015.0907.

58 Mao D, Zheng Y, Xu F, Han X, Zhao H. HMGB1 in nervous system diseases: A common biomarker and potential therapeutic target. Front Neurol 2022; 13: 1029891.

59 Gao T-L, Yuan X-T, Yang D, Dai H-L, Wang W-J, Peng X, et al. Expression of HMGB1 and RAGE in rat and human brains after traumatic brain injury. J Trauma Acute Care Surg 2012; 72: 643–9.

60 Qiu X, Guo Y, Liu M-F, Zhang B, Li J, Wei J-F, et al. Single-cell RNA-sequencing analysis reveals enhanced non-canonical neurotrophic factor signaling in the subacute phase of traumatic brain injury. CNS Neurosci Ther 2023; 29: 3446–59.

61 Tang Y-L, Fang L-J, Zhong L-Y, Jiang J, Dong X-Y, Feng Z. Hub genes and key pathways of traumatic brain injury: bioinformatics analysis and in vivo validation. Neural Regeneration Res 2020; 15: 2262.

62 Zhang Z, Yu J, Wang P, Lin L, Liu R, Zeng R, et al. iTRAQ-based proteomic profiling reveals protein alterations after traumatic brain injury and supports thyroxine as a potential treatment. Mol Brain 2021; 14: 1–21.

63 Gyoneva S, Ransohoff RM. Inflammatory reaction after traumatic brain injury: therapeutic potential of targeting cell–cell communication by chemokines. Trends Pharmacol Sci 2015; 36: 471–80.

64 Emerich DF, Skinner SJ, Borlongan CV, Vasconcellos AV, Thanos CG. The choroid plexus in the rise, fall and repair of the brain. Bioessays 2005; 27. DOI:10.1002/bies.20193.

65 Nave KA, Werner HB. Myelination of the nervous system: mechanisms and functions. Annu Rev Cell Dev Biol 2014; 30: 503–33.

66 Lee Y, Morrison BM, Li Y, Lengacher S, Farah MH, Hoffman PN, et al. Oligodendroglia metabolically support axons and contribute to neurodegeneration. Nature 2012; 487: 443–8.

67 Saab AS, Tzvetavona ID, Trevisiol A, Baltan S, Dibaj P, Kusch K, et al. Oligodendroglial NMDA Receptors Regulate Glucose Import and Axonal Energy Metabolism. Neuron 2016; 91: 119–32.

68 Mierzwa AJ, Marion CM, Sullivan GM, McDaniel DP, Armstrong RC. Components of myelin damage and repair in the progression of white matter pathology after mild traumatic brain injury. J Neuropathol Exp Neurol 2015; 74: 218–32.

69 Franklin RJM, Ffrench-Constant C. Regenerating CNS myelin - from mechanisms to experimental medicines. Nat Rev Neurosci 2017; 18: 753–69.

