## Supplementary Fig.S for "Thyroid Hormone T4 Mitigates Traumatic Brain Injury by Dynamically Remodeling Cell Type Specific Genes, Pathways, and Networks in Hippocampus and Frontal Cortex"

### Supplementary Materials Table of Contents

**Supplementary Fig.S1. Drop-seq library quality control (QC) features……………………………………2**

**Supplementary Fig.S2. Effects of TBI and T4 on frontal cortex and hippocampus neuron subtype cells across timepoints and conditions……………………………………………………………………………...3**

**Supplementary Fig.S3. Cell number distribution across the two tissues, timepoints and conditions………4**

**Supplementary Fig.S4. Top enriched pathways responsive to mTBI across timepoint and tissues………...5**

**Supplementary Fig.S5. Top enriched pathways responsive to T4 treatment across timepoints and tissues when only considering DEGs from T4 vs mTBI without comparing with DEGs in mTBI vs sham………..6**

### The following supplementary tables were uploaded separately as excel files:

Supplementary TableS1. List of differentially expressed genes (DEGs) after mTBI (TBI vs Sham) in the hippocampus and frontal cortex cell types.

Supplementary TableS2. List of differentially expressed genes (DEGs) after T4 treatment (T4 vs mTBI) identified in the hippocampus and frontal cortex cell types.

Supplementary TableS3. List of differentially expressed genes (DEGs) after mTBI (mTBI vs Sham) and T4 treatment (T4 vs mTBI) identified in the hippocampus and frontal cortex neuronal cell subtypes.

Supplementary TableS4. List of enriched pathways using differentially expressed genes (DEGs) identified after mTBI (mTBI vs Sham) in the hippocampus and frontal cortex cell types.

Supplementary TableS5. List of enriched pathways using differentially expressed genes (DEGs) identified after T4 treatment (T4 vs mTBI) in the hippocampus and frontal cortex cell types.

Supplementary TableS6. List of module genes, linear regression results, and enriched pathways identified in mTBI-specific modules for the cell type specific network analysis using SCING.

Supplementary TableS7. List of enriched human disease and traits using differentially expressed genes (DEGs) identified after mTBI (mTBI vs Sham) in the hippocampus and frontal cortex cell types.

Supplementary TableS8. List of enriched human disease and traits using differentially expressed genes (DEGs) identified after T4 treatment (T4 vs mTBI) in the hippocampus and frontal cortex cell types.

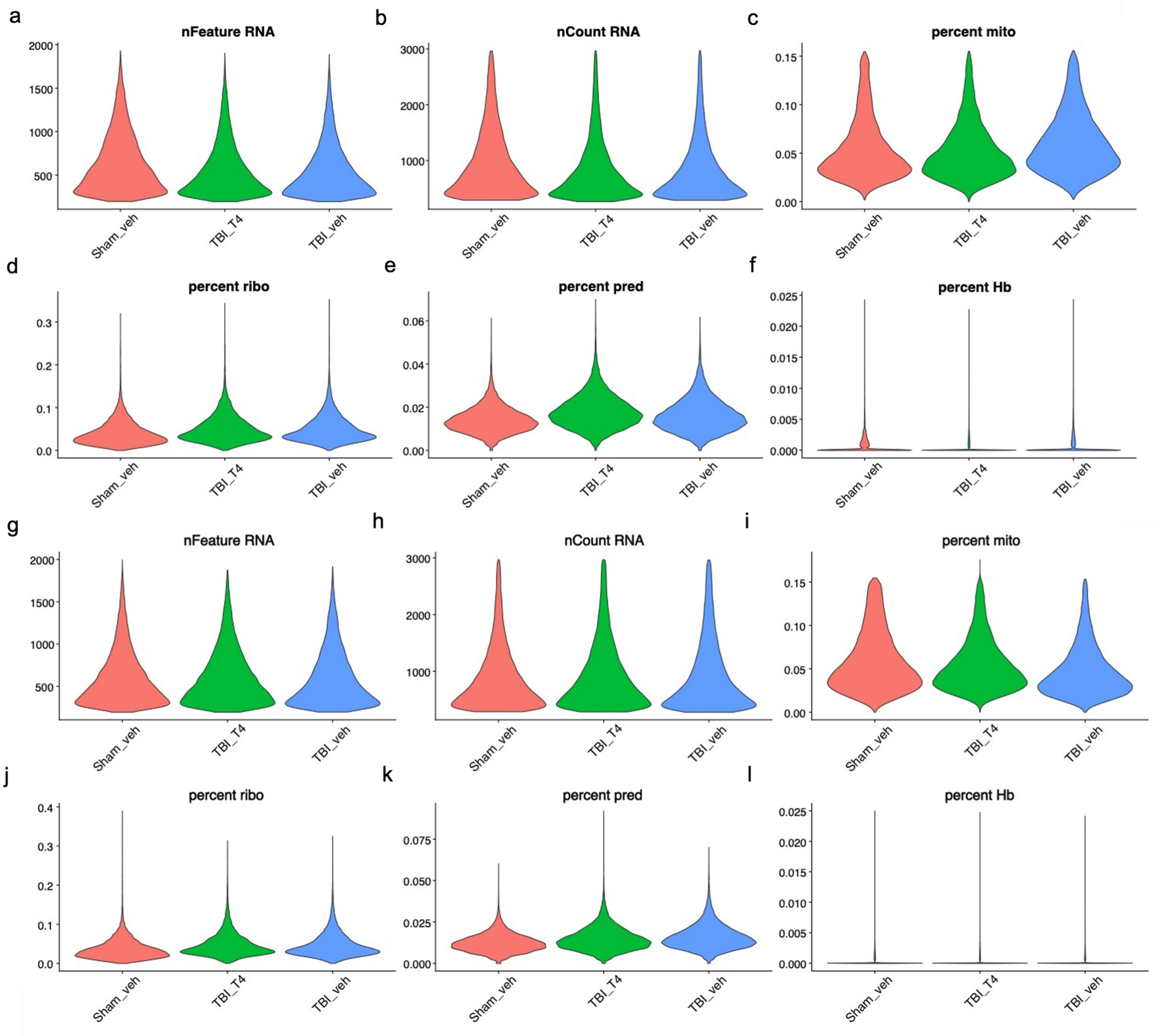


**Supplementary Fig.S1. Drop-seq library quality control (QC) features.** (a-f) Violin plot of the number of unique features (genes) detected (a), number of RNA counts (b), percent mitochondrial genes (c), percent ribosomal genes (d), percent of predicted genes (e) and percent hemoglobin genes (f) per single cell from the frontal cortex in each group. (g-l) Violin plot of the number of unique features (genes) detected (g), number of RNA counts h), percent mitochondrial genes (i), percent ribosomal genes (j), percent of predicted genes (k) and percent hemoglobin genes (l) per single cell from the hippocampus in each group.

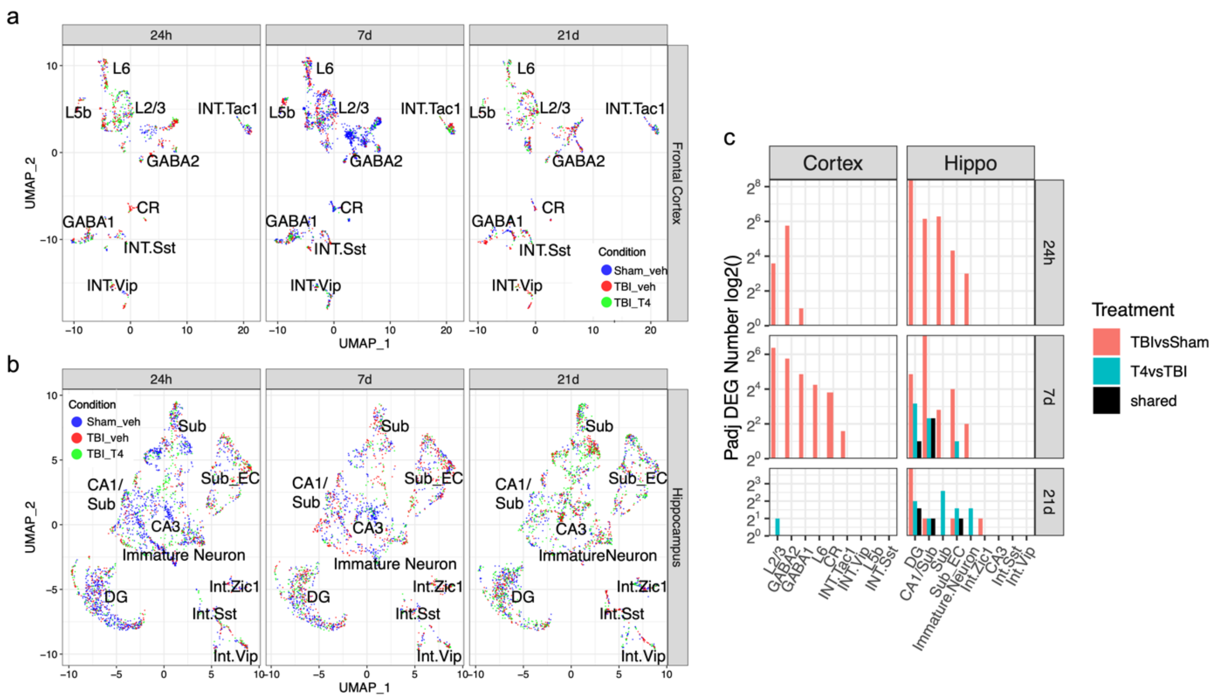

**Supplementary Fig.S2. Effects of TBI and T4 on frontal cortex and hippocampus neuron subtype cells across timepoints and conditions.** (a and b) UMAP plot of frontal cortex (a) and hippocampus (b) neuron subtypes across three timepoints (24h, 7d and 21d) with each cell colored according to condition (Sham veh, TBI veh and TBI T4). (c) Bar plot of the number of differentially expressed genes (DEGs) identified in frontal cortex and hippocampus neuronal subtype cells.

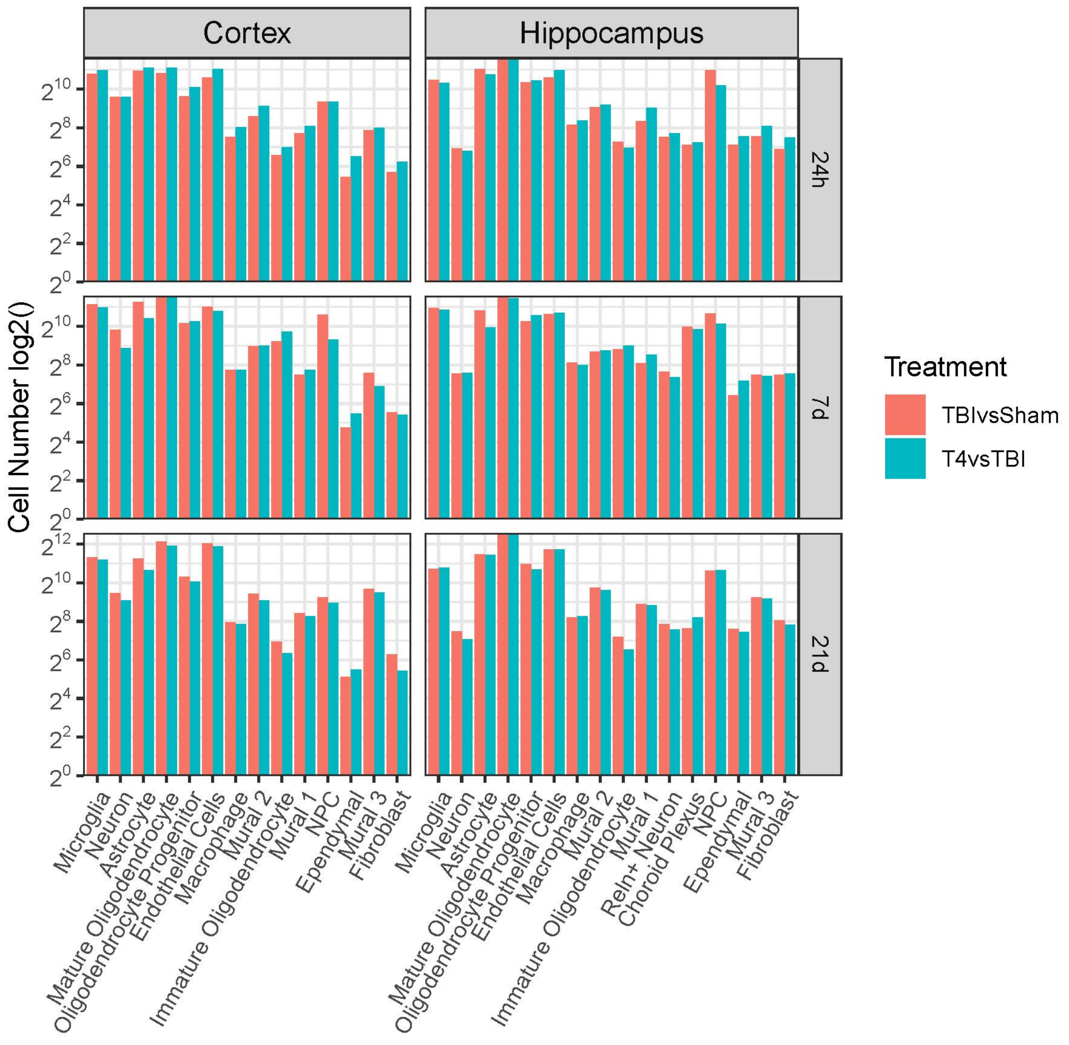


**Supplementary Fig.S3. Cell number distribution across the two tissues, timepoints and conditions.** Bar plot comparing the cell numbers across the three treatments and three timepoints in the different cell types identified in the hippocampus and cortex region.

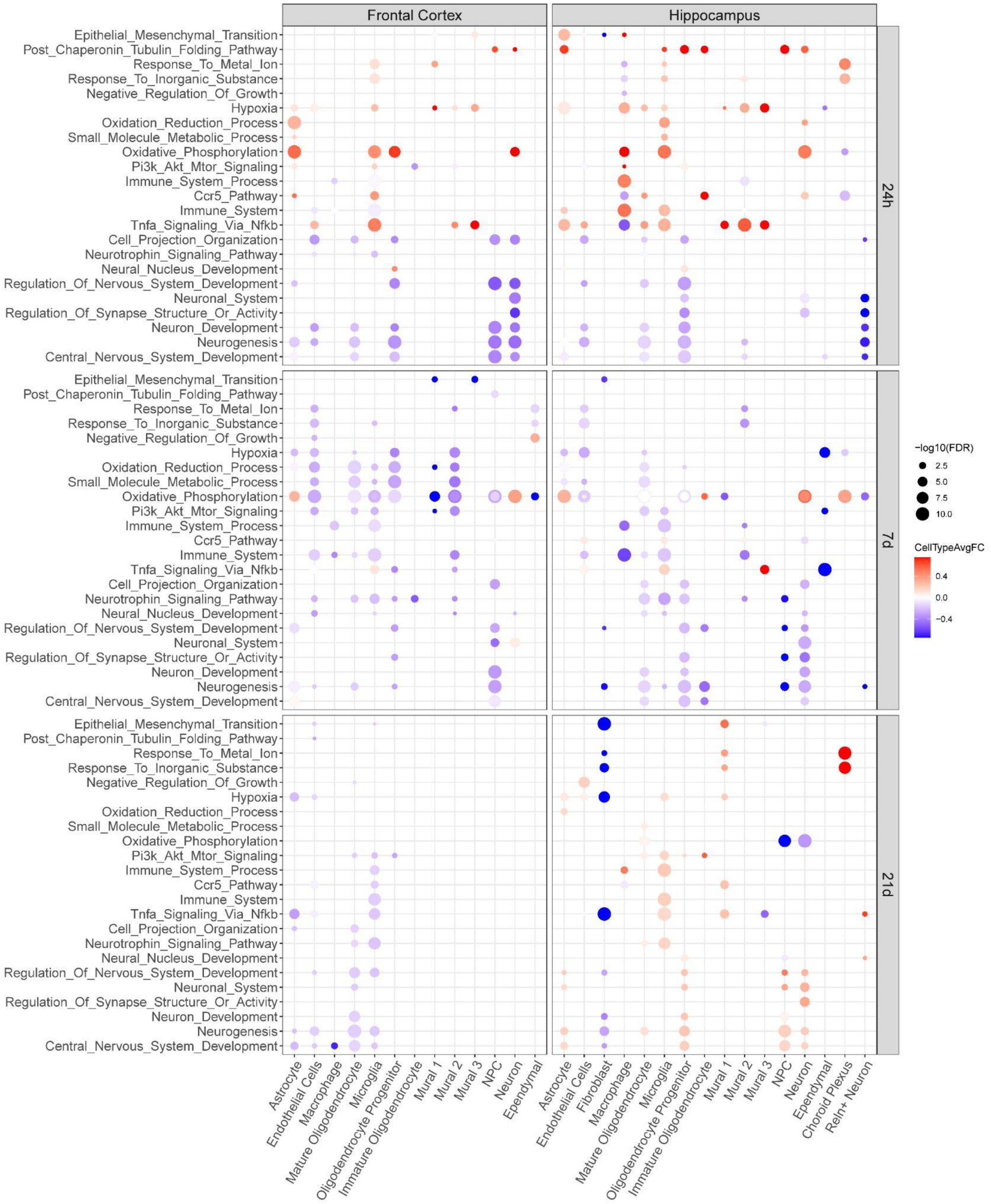


**Supplementary Fig.S4. Top enriched pathways responsive to mTBI across timepoint and tissues.** Each dot is colored by the average log fold change between mTBI and sham control cells within that cell type for the significant DEGs which overlap the indicated pathway. The size of each dot is proportional to the statistical enrichment significance in the form of −log10(FDR). Cell types and pathways have been clustered with hierarchical clustering.

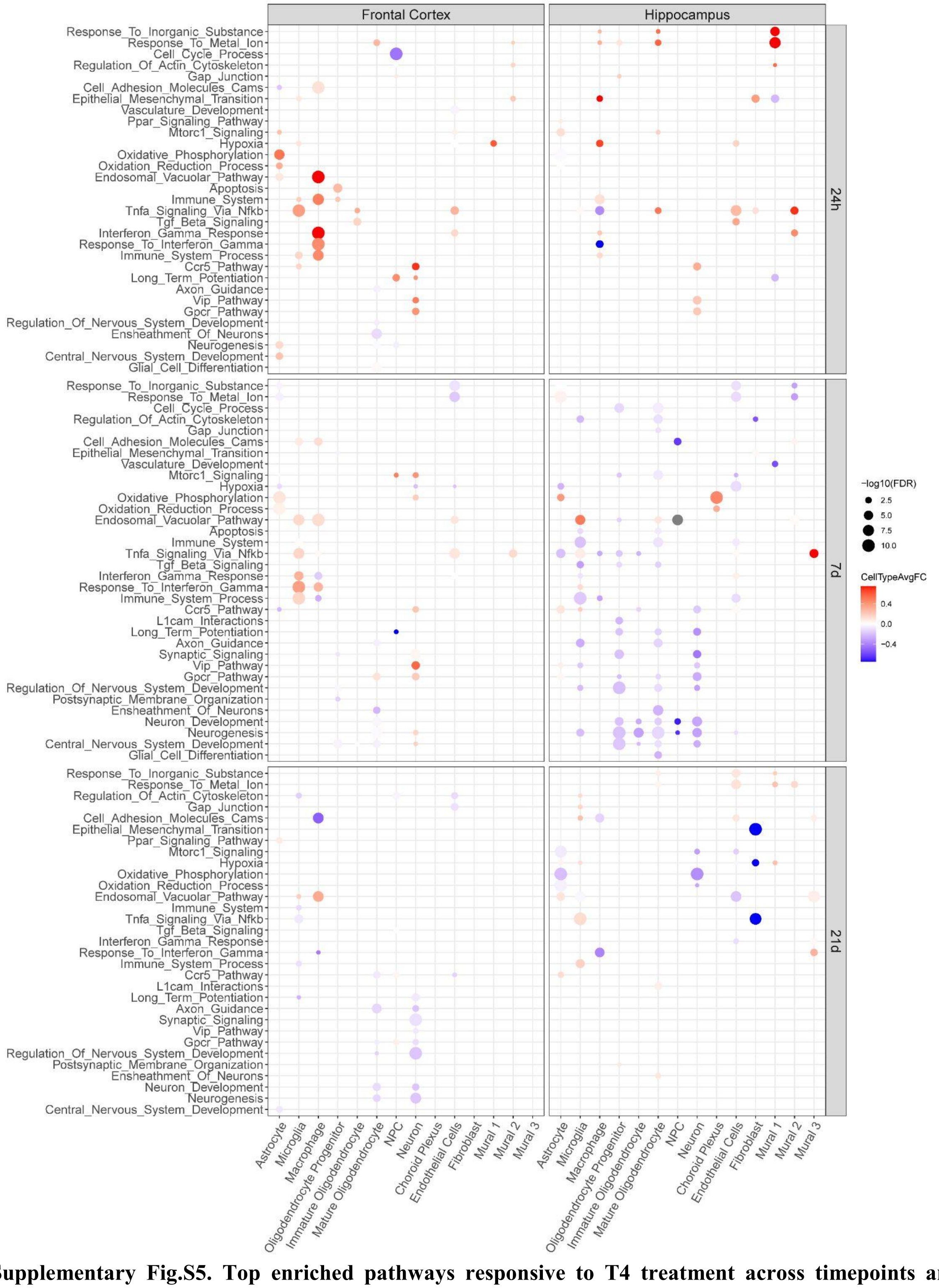


**Supplementary Fig.S5. Top enriched pathways responsive to T4 treatment across timepoints and tissues when only considering DEGs from T4 vs mTBI without comparing with DEGs in mTBI vs sham.** Each dot is colored by the average log fold change between mTBI and sham control cells within that cell type for the significant DEGs which overlap the indicated pathway. The size of each dot is proportional to the statistical enrichment significance in the form of −log10(FDR). Cell types and pathways have been clustered with hierarchical clustering.
